# Multiple QTL mapping in autopolyploids: a random-effect model approach with application in a hexaploid sweetpotato full-sib population

**DOI:** 10.1101/622951

**Authors:** Guilherme da Silva Pereira, Dorcus C. Gemenet, Marcelo Mollinari, Bode A. Olukolu, Joshua C. Wood, Federico Diaz, Veronica Mosquera, Wolfgang J. Gruneberg, Awais Khan, C. Robin Buell, G. Craig Yencho, Zhao-Bang Zeng

## Abstract

In developing countries, the sweetpotato, *Ipomoea batatas* (L.) Lam. (2*n* = 6*x* = 90), is an important autopolyploid species, both socially and economically. However, quantitative trait loci (QTL) mapping has remained limited due to its genetic complexity. Current fixed-effect models can only fit a single QTL and are generally hard to interpret. Here we report the use of a random-effect model approach to map multiple QTL based on score statistics in a sweetpotato bi-parental population (‘Beauregard’ *×* ‘Tanzania’) with 315 full-sibs. Phenotypic data were collected for eight yield component traits in six environments in Peru, and jointly predicted means were obtained using mixed-effect models. An integrated linkage map consisting of 30,684 markers distributed along 15 linkage groups (LGs) was used to obtain the genotype conditional probabilities of putative QTL at every cM position. Multiple interval mapping was performed using our R package QTLPOLY and detected a total of 41 QTL, ranging from one to ten QTL per trait. Some regions, such as those on LGs 3 and 15, were consistently detected among root number and yield traits and provided basis for candidate gene search. In addition, some QTL were found to affect commercial and noncommercial root traits distinctly. Further best linear unbiased predictions allowed us to characterize additive allele effects as well as to compute QTL-based breeding values for selection. Together with quantitative genotyping and its appropriate usage in linkage analyses, this QTL mapping methodology will facilitate the use of genomic tools in sweetpotato breeding as well as in other autopolyploids.

---

Genetic analyses in polyploid species pose extra challenges in comparison to diploid species, in spite of the evolutionary benefits that duplication of whole sets of chromosomes may have brought (Comai 2005; Van De Peer *et al*. 2009). When it comes to molecular markers, a co-dominant, biallelic single nucleotide polymorphism (SNP) directly informs on the genotypes of a diploid locus, but the best it can do alone in a polyploid locus is to inform on its allele dosage. In diploid species, molecular markers are usually qualitatively scored, and there are several methodologies and tools for performing analysis on genetic linkage (e.g., Stam 1993; Margarido *et al*. 2007) and quantitative trait loci (QTL) mapping (e.g., Broman *et al*. 2003; Da Costa E Silva *et al*. 2012a). In allopolyploid species, such as cotton (Wu *et al*. 2015) and wheat (Hulse-Kemp *et al*. 2015), where preferential paring dictates meiotic chromosome behavior much like diploids, existing approaches can be readily applied. However, despite many successful studies in diploids and allopolyploids, QTL mapping in autopolyploids remains difficult. In fact, unlike diploid mapping populations, which can have two to four segregating QTL genotypes (in case of inbred or outbred species, respectively), autopolyploid mapping populations can have a much wider range of possible genotypes per locus. For example, there are up to 36, 400 or 4,900 possible genotypes from crosses between two tetra-, hexa-or octoploid outbred parents, respectively.

Single-dose markers, segregating in 1:1, 3:1 or 1:2:1 fashion, have limited information for building integrated genetic maps in autopolyploids, and can only be used for developing separate parental maps (Shirasawa *et al*. 2017) or rather than the desired integrated linkage groups (Balsalobre *et al*. 2017). In order to make use of multiple-dose markers, the first step is to perform dosage or quantitative SNP calling. Although most methods were designed for tetraploid species (e.g., Voorrips *et al*. 2011; Schmitz Carley *et al*. 2017), additional studies have tackled this problem, and methods including higher ploidy levels are now available (Serang *et al*. 2012; Gerard *et al*. 2018). For building integrated genetic maps, tetraploid species can use the well-established TETRA PLOIDSNPMAP (Hackett *et al*. 2016). For higher ploidy species, MAPPOLY (Mollinari and Garcia 2018) is a better option than POLYMAPR (Bourke *et al*. 2018), because the latter is limited to tetra- and hexaploid species, and lacks the ability to robustly map all multiple-dose markers based on hidden Markov models (HMM). With an integrated map, one can calculate the putative QTL genotype conditional probabilities, ideally using appropriate HMM (Hackett *et al*. 2016; Mollinari and Garcia 2018). Based on a model in Kempthorne (1955), an interval mapping (IM) method has been proposed as a first approach to map QTL in autotetraploids in a form of a regression weighted by the conditional probabilities (Hackett *et al*. 2001), which also turned out to be expanded for an autohexaploid species (van Geest *et al*. 2017).

For a general ploidy level *m*, this single-QTL model can be written as

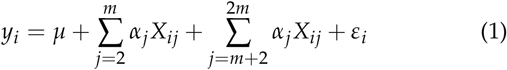

where *y*_*i*_ is the phenotypic value of individual *i, µ* is the intercept, *α*_*j*_ is the additive effect of allele *j, X*_*ij*_ is the conditional probability of allele *j* in individual *i*, and *ε*_*i*_ is the residual error. The constraints *α*_1_ = 0 and *α*_*m*+1_ = 0 are naturally imposed to satisfy the conditions 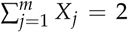 and 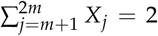, so that *µ* is a constant hard to interpret due to these constraints (Hackett *et al*. 2001). Notice that 2*m −* 2 effects need to be estimated, i.e. tetra-, hexa- or octoploid models will have six, ten or 14 main effects, respectively. In order to answer the key question of whether the additive allele effects are different from zero (the null hypothesis), likelihood-ratio tests (LRT) are performed along positions on a genetic map. Commonly, the tests are presented as “logarithm of the odds” (LOD scores), where 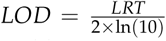. In order to declare a QTL, empirical LOD thresholds are computed for each trait using permutations (Churchill and Doerge 1994). As the only current solution, this approach has been widely used so far (e.g., van Geest *et al*. 2017; Schumann *et al*. 2017; Massa *et al*. 2018). However, limitations in fitting multiple-QTL models have been presented mostly due to the possibility of over-parameterization or the lack of optimized algorithms for model selection (Mengist *et al*. 2018; Klaassen *et al*. 2019).

Variance component methods have been used for performing QTL mapping in related individuals of complex population structures or families in humans (Lippert *et al*. 2014), animals (Druet *et al*. 2008) and plants (Crepieux *et al*. 2005). In common, these approaches take into account the flexibility of mixed models in dealing with the correlated QTL effects among individuals due to shared alleles identical-by-descent (IBD) by each relative pair at a particular location in the genome. For bi-parental populations of autopolyploids, although allele effects are usually regarded as fixed, the QTL genotype effects can be treated as random. Since a higher ploidy level leads to a much larger number of allele combinations, genotypic effects may be very hard to assess from the small population sample sizes usually available. In this case, the integrated genetic map provides key information on the inheritance of chromosomal segments from parents to progeny (Mollinari and Garcia 2018), making up the basis for IBD allele sharing estimations. If a locus is linked to a region underlying the variation of a trait of interest, higher IBD allele sharing for that locus is expected among individuals with similar phenotypic values (Almasy and Blangero 2010). Thus, the key parameter in this model are the variance components attributable to putative QTL, that determines the presence of linkage. Because only one parameter per QTL (the variance component) needs to be estimated, one could try to build a multiple-QTL model for polyploids, inspired by the corresponding multiple interval mapping (MIM) for diploid mapping populations (Kao *et al*. 1999), without the risk of model over-parameterization.

A multiple-QTL mapping approach may benefit several autopolyploid horticultural (e.g, potato, blueberry, kiwifruit, strawberry), ornamental (e.g., rose, chrysanthemum), forage (e.g., alfalfa, guinea grass) and field (e.g., sugarcane) crops. The sweet-potato [*Ipomoea batatas* (L.) Lam. (2*n* = 6*x* = 90)] is a staple food in several developing countries, with a production of 112 million tons worldwide in 2017 (FAO 2019). Particularly, it has attracted growing interest due to its characteristics for food and nutrition security (Mwanga *et al*. 2017). In addition to carbohydrates, dietary fiber, vitamins and minerals, orange-fleshed sweetpotatoes provide high levels of *β*-carotene to fight vitamin A deficiency in vulnerable populations, such as those in sub-Saharan Africa (Low *et al*. 2017). In order to increase production and meet farmer’s and market needs, it is imperative to make molecular-assisted selection an effective part of sweet-potato breeding programs. Toward this end, one of the first steps is characterizing the genetic architecture of traits of interest, such as those related to storage root yield and quality, and resistance to biotic and abiotic stresses (Khan *et al*. 2016). In spite of being considered an “orphan” crop, there have been recent advances in building genome references from its wild diploid relatives (Wu *et al*. 2018), optimizing a genotyping-by-sequencing protocol (GBSpoly) for high-throughput SNP genotyping (Wadl *et al*. 2018), and building a high-density integrated genetic map (Mollinari *et al*. 2019, in preparation). In this paper, we introduce a random-effect multiple interval mapping (REMIM) model for autopolyploids. Using a genome-assisted, GBSpoly-based integrated genetic map from a sweetpotato bi-parental population, we map QTL for yield-related traits with our open-source software, QTLPOLY.

## Materials and Methods

### Full-sib Population

A bi-parental mapping population (named BT) comprising 315 individuals was developed by crossing an orange-fleshed American variety, ‘Beauregard’ (CIP440132), and a non-orange-fleshed African landrace, ‘Tanzania’ (CIP440166), as male and female parents, respectively. The parents show contrasting phenotypes for several traits such as dry matter, *β*-carotene and sugar con-tent, and susceptibility to biotic (e.g., virus disease) and abiotic (e.g., drought) stresses. ‘Beauregard’ is known as to have higher yield than ‘Tanzania’, and the current QTL mapping study will focus on the yield components.

### Phenotypic Analyses

#### Field trials

In addition to the 315 full-sibs, parents (each replicated twice) and another variety, ‘Daga’ (CIP199062.1), were used as checks in order to make up a total of 320 individuals per replication in an 80 × 4 alpha-lattice design. Virus-free planting material derived from tissue culture was obtained from the CIP-Peru Genebank in La Molina. The clones were grown in a screen house in CIP sub-station San Ramon, and the planting material multiplied under low-disease pressure field conditions in Satipo, where cuttings for the six experiments were obtained. Four experiments were conducted in Ica (14°01’ S and 75°44’ W, 420 m), with two independent trials over two seasons, and one experiment each was conducted in San Ramon (11°07’ S and 75°21’ W, 828 m) and Pucallpa (8°23’ S and 74°31’ W, 154 m). The number of replications were two at Ica and three at San Ramon and Pucallpa. In all trials, 1 m and 0.3 m of inter- and intra-row spacing was used, respectively. In the first season at Ica (from February 25 to June 29, 2016), the plot size was 6 m^2^ of 16 plants arranged in four rows (four plants per row) with one empty row between plots. In the second season at Ica (from November 15 2016 to March 17, 2017), the plot size was 4.8 m^2^ of 16 plants arranged in two rows (eight plants per row) with no empty row between plots. In San Ramon (from May 14 to September 15, 2016) and Pucallpa (from July 1 to November 4, 2016), the plot size was 9 m^2^ of 30 plants arranged in three rows (ten plants per row) with no empty row between plots.

#### Phenotypic data

Eight yield-related phenotypes were collected per plot at harvest, *∼* 120 days after transplanting (see File S1). For analysis purposes, foliage and root yield data were standardized by plot size (relative to the largest) and converted to tons per hectare (t · ha^*-*1^) to allow comparisons across trials. Number of roots was divided by the number of plants in the plot. The total number of storage roots per plant (TNR) and total root yield (RYTHA) considered all storage roots from the whole plot regardless of their individual weight. Number of commercial roots per plant (NOCR) and commercial root yield (CYTHA) considered only storage roots of marketable size (*≥*100 g for African market). Number of noncommercial roots per plant (NONC) and noncommercial root weight (NCYTHA) were obtained from the difference between total and commercial roots. Foliage yield (FYTHA) was measured by weighing all above-ground biomass per plot. Finally, commercial index (CI) was calculated as the ratio between CYTHA and total biomass (i.e. the sum of RYTHA and FYTHA).

#### Multi-environment phenotypic model

We considered each one of the six field trials as an environment. Jointly predicted means for each full-sib were obtained by using the following mixed-effect model

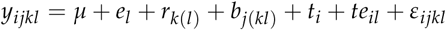

where *y*_*ijkl*_ is the phenotype of the *i*^th^ treatment in the *j*^th^ block within the *k*^th^ replicate at the *l*^th^ environment, *µ* is the overallmean, *e*_*l*_ is the fixed effect of the *l* ^th^ environment (*l* = 1, …, *L*; *L* = 6), *r*_*k*(*l*)_ is the fixed effect of the *k*^th^ replicate (*k* = 1, …, *K*; *K* = 2 or 3 depending on the environment) at the *l*^th^ environment, *b*_*j*(*kl*)_ is the random effect of the *j*^th^ block (*j* = 1, …, *J*; *J* = 80) within the *k*^th^ replicate at the *l*^th^ environment with 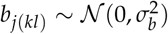, *t*_*i*_ is the effect of the *i*^th^ treatment (*i* = 1, …, *I*; *I* = 318), *te*_*il*_ is the effect of treatment by environment interaction, and *ε*_*ijkl*_ is the random residual error with 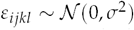. The treatment effect (*t*_*i*_) was separated into two groups, in which *g*_*i*_ is the random effect of the *i*^th^ genotype (*i* = 1, …, *I*_*g*_; *I*_*g*_ = 315) with 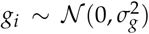, and *c*_*i*_ is the fixed effect of the *i*^th^ check (*i* = *I*_*g*_ + 1, …, *I*_*g*_ + *I*_*c*_; *I*_*c*_ = 3). Similarly, treatment by environment interaction (*te*_*il*_) was separated into the random effect of genotype by environment interaction (*ge*_*il*_) with 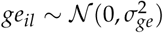, and the fixed effect of check by environment interaction (*ce*_*il*_). We removed the check by environment interaction effect (*ce*_*il*_) from the model if Wald’s test was not significant (*p <* 0.01). Variance components were estimated by restricted maximum likelihood (REML) using GENSTAT (v16; VSN International 2014). Mean-basis broad-sense heritabilities (*H*^2^) were calculated as the ratio between genotypic and phenotypic variances as

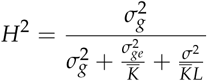

where 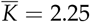 is the harmonic mean of the number of replicates across environments. The R package PSYCH (v1.8.10; Revelle 2018) was used to calculate and plot Pearson’s correlations (significance \**p <* 0.05, \*\**p <* 0.01 and \*\*\**p <* 0.001) among the individual predicted means.

### Genotypic Analyses

#### GBSpoly and dosage calling

A modified GBS protocol called GBSpoly was carried out according to Wadl *et al*. (2018) and described in detail for BT population by (Mollinari *et al*. 2019, in preparation). In brief, total DNA was extracted and double restricted using *Cvi*AII-*Tse*I enzyme combination for all fullsibs and parents (each parent replicated 10 times). Restriction fragments were ligated to adapters, size selected and amplified. Adapters contained an 8-bp buffer sequence in addition to sample-specific variable length barcodes (6-9 bp). Each 64-plex library was sequenced using eight lanes of Illumina HiSeq 2500 system in order to ensure optimal read depth for dosage calling. We trimmed the 8-bp buffer sequence from the reads using the FASTX-TOOLKIT (available at hannonlab.cshl.edu/fastx_toolkit/). A modified version of TASSEL-GBS pipeline (v4.3.8), called TASSEL4-POLY (Pereira *et al*. 2018, available at https://github.com/gramarga/tassel4-poly) was used to demultiplex and to count and store the actual read depth for all loci in variant call format (VCF) files (see File S2). We used BOWTIE2 (Langmead and Salzberg 2013) to align 64-bp tags against the *I*. *trifida* and *I*. *triloba* genomes, two sweetpotato wild relative diploid species (Wu *et al*. 2018, available at http://sweetpotato.plantbiology.msu.edu). Finally, the software SUPERMASSA (Serang *et al*. 2012, available at https://bitbucket.org/orserang/supermassa) was used to perform multi-threading dosage call through a wrapper function named VCF2SM (Pereira *et al*. 2018, available at https://github.com/gramarga/vcf2sm).

#### Linkage mapping

A linkage map was constructed by (Mollinari *et al*. 2019, in preparation) using the R package MAPPOLY (Mollinari and Garcia 2018, available at https://github.com/mmollina/ mappoly) (see File S3). In brief, we computed two-point recombination fractions between all 38,701 non-redundant, high quality GBSpoly-based markers, and sorted the most likely linkage phase between each marker pair. Markers were then grouped into 15 linkage groups (LGs) by using the Unweighted Pair Group Method with Arithmetic Mean (UPGMA) hierarchical clustering method. For each LG, markers were first ordered using multidimensional scaling as implemented in the R package MDSMAP (Preedy and Hackett 2016), and then local order was refined based on the reference genomes (Wu *et al*. 2018). Finally, map distances were re-estimate individual posterior probabilities from SUPERMASSA dosage calls. The final integrated, completely phased map was composed of 30,684 markers distributed along 15 LGs with a total length of 2,708.4 centiMorgans (cM) and no major gaps between markers (11.35 markers every cM, on average). Multi-point genotype conditional probabilities of putative QTL were estimated for every individual given the final map using an HMM algorithm (Lander and Green 1987; Jiang and Zeng 1997) adapted for polyploids (Mollinari *et al*. 2019, in preparation) as implemented in MAPPOLY. Since 17 full-sibs were filtered out along the map construction (Mollinari *et al*. 2019, in preparation), only the remaining 298 individuals were ultimately used for QTL mapping.

#### QTL Mapping Analyses

Under some fairly conventional assumptions (exclusive bivalent formation, no preferential pairing, and no double reduction), an autopolyploid individual of a species with an even ploidy level *m* can produce up to 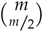 or “*m* choose *m*/2” different gametes with the same probability. As an example, consider two contrasting parents, A and B, of a hexaploid species (such as sweetpotato) and their respective genotypes for a QTL as *abcde f* and *ghijkl*, each one with potentially six different alleles. Under the previous assumptions, each parent can produce up to 20 different gametes. Therefore, the cross A *×* B would generate up to 400 possible different genotypes. We will use the number of gametes and genotypes of a hexaploid species to define vector and matrix dimensions in the QTL mapping model from now on. Obviously, the model can be easily adapted to any polyploid species with an even ploidy level by simply changing these dimensions accordingly.

#### REMIM model and hypothesis testing

Taking a full-sib population with *n* individuals derived from a cross between two hexaploid parents, A and B, the multiple-QTL mapping model is expressed by

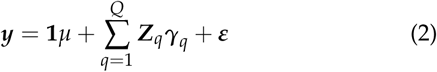

where ***y*** is the *n ×* 1 vector of phenotypic values, *µ* is the fixed effect of population mean, ***γ***_*q*_ is the 400 *×* 1 random vector of genotype effects of QTL *q* (*q* = 1, …, *Q*) with 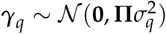, and ***ε*** is the *n ×* 1 random vector of residual error with ***ε*** ∼*𝒩*(**0, *I****σ*^2^). **1** and ***I*** are an *n*× 1 vector of 1’s and an *n* ×*n* identity matrix, respectively, ***Z***_*q*_ is the *n* × 400 incidence matrix of genotype conditional probabilities of QTL *q*, and **Π** is a 400 × 400 matrix of proportion of shared alleles IBD between the 400 possible genotypes. These IBD allele sharing proportions range from zero (no shared alleles) to one (six shared alleles) and relate to the additive effects of within-parent alleles. For a full-sib progeny, the IBD expected value is 0.5.

Assuming that the random-effect QTL are uncorrelated, each with expectation zero, the expectation of the vector of phenotypic values ***y*** is

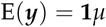

and its variance-covariance matrix is

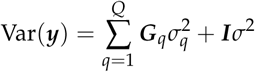

where 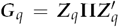 is the *n* × *n* additive relationship matrix between all *n* full-sibs on the putative QTL *q*. Here, our interest is in testing

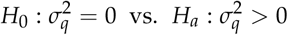

i.e., whether QTL *q* contributes to the variation in ***y*** or not, so that several tests have to be performed along the genome. As part of the algorithm described next, we test for the presence of multiple QTL in consecutive rounds. In practice, we compute and store a ***G***_*q*_ matrix for every putative QTL *q*, representing genomic positions at a certain step size (e.g., every 1 cM). In this case, the Model 2 can be rewritten as

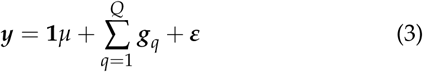

where ***g***_*q*_ is an *n ×* 1 random vector of the individual effects for the QTL *q* with 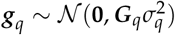.

We compute linear score statistics according to Qu *et al*. (2013) at every position and compare its *p*-value to a prescribed critical value. The *p*-values are continuous over the unit interval as a result of weighted sums of the scores from the profiled likelihood. The test is exact (nonasymptotic) when there is only one QTL, while a moment-based approximation to the null distribution is used when two or more QTL are present in the model (Qu *et al*. 2013). Herein, we conveniently take the “logarithm of *p*” as *LOP* = *-* log_10_(*p*) for graphic representation and supporting interval calculation purposes. Support intervals are defined as the QTL peak neighboring region with *LOP* greater than or equal *LOP - d*, where *d* is a constant which subtracts the highest *LOP* (thus from the QTL peak) in that region, as similarly proposed for the statistic LOD scores (Lander and Green 1987).

#### QTL detection and characterization

In order to select QTL, we adapted the MIM methodology described by Kao *et al*. (1999) to a random-effect model framework as follows:

1. *Forward search* adds one QTL at a time to the model at the position with the highest score statistic if the *p*-value is smaller than a pointwise significance threshold level (e.g., *p <* 0.01), and fits it into the model. Consecutive rounds are carried out conditioning the search of a new QTL to the one(s) in the model until no more positions can reach the threshold. A window size (e.g., of 15 cM) is avoided on either side of QTL already in the model when searching for a new QTL;
2. *Model optimization* follows rounds of position refinement and backward elimination when no more QTL can be added in the forward search step. In turns, a QTL position is updated conditional to all the other QTL in the model, and its score statistic is reevaluated at a more stringent significance threshold level (e.g., *p <* 10^*-*3^ or 10^*-*4^), when the QTL may be dropped. The final set of QTL is defined when all selected positions are significant and, thus, no more positions change or QTL are dropped;
3. Forward search (now with a threshold value as stringent as the one used for backward elimination, e.g., *p <* 10^*-*3^ or 10^*-*4^) as well as model optimization procedures are repeated until no more QTL are added (via forward search) or dropped (via backward elimination). Finally, *QTL profiling* is performed with the remaining significant QTL after the last round of model optimization has been carried out. The score statistics and their associated *p*-values are computed for all genomic positions conditional to the final set of QTL.

Notice that, as part of the strategy for selecting QTL, we were less stringent during the first step of *forward search*, so that we were able to allow more positions to be tested again during *model optimization*. In fact, power for detecting significant positions is expected to increase when conditioning the forward search as well as the backward elimination to other QTL already in the model (Da Costa E Silva *et al*. 2012a). For the forward search performed after the first backward elimination, we used the last threshold set from the backward elimination in order to avoid false positives.

Once the QTL were selected, we were able to estimate their variance components and compute QTL heritabilities, 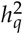, as the ratio between the QTL variance component and total variance. Given the parameter estimates, QTL-based breeding values are directly obtained as the best linear unbiased predictions (BLUPs) of the QTL genotypes (i.e.,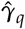) from Model 2. BLUPs of the 400 possible genotypes were further decomposed in order to estimate the additive allele effects (i.e., 6 for each parent as {*a*, …, *f*} and {*g*, …, *l*}) as well as the additive allele combination effects among two (i.e., 15 combinations for each parent as {*ab*, …, *cd*} and {*e f*, …, *kl*}), and three (i.e., 20 combinations for each parent as {*abc*, …, *def*} and {*ghi*, …, *jkl*}) alleles (Kempthorne 1955). Notice that in an F_1_ population, we can only study QTL that are different in alleles within the parents, not between. Also, due to the model assumptions of zero mean for random effects, allele and allele combination effects sum up to zero. These effects should be interpreted as the heritable contributions from parent to offspring, hence providing straightforward estimation of QTL-based breeding values to be used for selection.

### Simulations

We examined the performance of REMIM with 1,000 simulated quantitative traits with three QTL each. The QTL heritabilities were simulated as 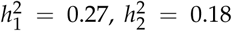 and 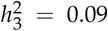 following their respective QTL genotype effect distributions as ***g***1 ∼ 𝒩 (**0, *G***_1_0.6), ***g***2 ∼ 𝒩 (**0, *G***_2_0.4) and ***g***3 ∼ 𝒩 (**0, *G***_3_0.2). The environmental error was simulated from a standard normal distribution, i.e. ***ε*** ∼ 𝒩 (**0, *I***1), while the population mean was simulated as zero, i.e. *µ* = 0. The QTL were randomly assigned to the BT population linkage map (*n* = 298), but no closer than 15 cM (our window size) from each other. One round of *forward search* followed by *model optimization* (steps 1 and 2 from the algorithm described above) was carried out for each simulated trait with combinations of different forward (0.01, 0.02 and 0.05) and backward (10^−2^, 10^−3^, 10^−4^ and 10^−5^) pointwise significance *p*-value thresholds. For comparison, we ran the fixed-effect interval mapping (FEIM, Model 1) with the same simulated traits using different genome-wide significance levels of (0.20, 0.15, 0.10 and 0.05) based on 1,000 permutations as LOD thresholds to declare significant QTLs (Churchill and Doerge 1994). The same step size of 2 cM was used in both approaches. *LOP* − *d* (from REMIM) and *LOD* − *d* (from FEIM) support intervals were calculated for three different *d* values (1.0, 1.5 and 2.0).

Following the definitions and summary statistics from Da Costa E Silva *et al*. (2012b), all QTL kept after the model optimization were considered “mapped”. A mapped QTL was considered “paired” (true QTL) if less than 15 cM apart from the simulated position, and a paired QTL was considered “matched” if included within a support interval of a mapped QTL. Finally, a mapped QTL was considered “mismatched” (false QTL) if it was not matched. We summarized detection power as the ratio between the number of paired QTL over the total number of simulated QTL, and the absolute distance differences between simulated and mapped positions were averaged out. Genomewide type I error or false discovery rate (FDR) was estimated for each support interval as the ratio between the number of mismatched QTL over the total number of mapped QTL. Finally, the proportion of matched QTL (coverage) as an approximation of support intervals was provided for each *d* value.

### Software implementation

We implemented the algorithm for detection and characterization of multiple QTL based on REMIM model in an R package called QTLPOLY (available at https://github.com/guilherme-pereira/qtlpoly). We integrated functions from the R package VARCOMP (v0.2-0; Qu *et al*. 2013) to compute the score statistics. The rounds of QTL search and model optimization use the variance components estimated in the previous round, so that the new estimates iterate faster. In addition, calculations for different genomic positions were paralleled in order to speed up the process by using the R base package PARALLEL (v3.5.2; R Core Team 2018). Final models were fitted using the R package SOMMER (v3.6; Covarrubias-Pazaran 2016), from which BLUPs were extracted and used for estimation of allele effects and QTL-based breeding values. Both VARCOMP and SOMMER packages use REML estimation to compute the variance components from the random-effect QTL model. Functions for plotting QTL profiles, effects and support intervals were based on GG-PLOT2 (v3.1.0; Wickham 2016). Additional functions for running FEIM model and multi-threaded permutations were included in QTLPOLY and were based on the `lm()` function from R base package STATS (v3.5.2; R Core Team 2018).

### Gene expression profiling

A developmental time-course of expression profiling data of ‘Beauregard’ was reported previously (Wu *et al*. 2018) and used with a parallel time-series of development with ‘Tanzania’ roots (Gemenet *et al*. 2019, submitted). In brief, ‘Beauregard’ and ‘Tanzania’ roots were harvested from four biological replicates at 10, 20, 30, 40 and 50 days after transplanting (DAT), and classified at 30, 40, and 50 DAT into fibrous and storage roots based on diameter as described by Wu *et al*. (2018). RNA and RNA-sequencing (RNA-seq) from ‘Tanzania’ were generated in parallel with the ‘Beauregard’ samples as described in Wu *et al*. (2018). All reads were cleaned, aligned to the *I*. *trifida* genome (Wu *et al*. 2018), and fragments per kilobase exon model per million mapped reads (FPKM) determined as described previously in Lau *et al*. (2018) with the one exception that the ‘Tanzania’ 30 DAT storage root sample was sub-sampled for 30 million reads. To provide a comparison of expression abundances in the roots to leaves, ‘Beauregard’ and ‘Tanzania’ plants were grown as described in Lau *et al*. (2018) for control conditions and RNA-seq libraries from leaves processed as described above. For the final FPKM matrix, genes encoded by the chloroplast were removed.

### Data Availability

Raw sequence reads are available at NCBI, under BioProject numbers PRJNAXXXXXX (DNA), and PRJNA491292 and PRJ-NAXXXXXX (RNA) [*to be released upon publication*]. The expression abundance matrix is available at Dryad Digital Repository [*to be released upon publication*]. Remaining supplemental files are available at FigShare [*to be released upon publication*]. File S1 contains phenotypic data. File S2 contains VCF files. File S3 contains genetic map information. QTLPOLY software used for QTL mapping analyses and simulations is available at GitHub (https://github.com/guilherme-pereira/qtlpoly).

## Results

### Trait heritabilities and correlations

Each one of the eight yield-related traits from six environments were analyzed using a multi-environment mixed-effect model, from which we were able to obtain jointly predicted means for each full-sib and variance component estimates (Table 1). Parents showed contrasting means for all traits, with ‘Beauregard’ presenting higher means for number of roots and root yield (both commercial and non-commercial) and commercial index when compared to ‘Tanzania’, which surpassed ‘Beauregard’ only for foliage yield. Interestingly, transgressive segregation was observed among the full-sibs for all traits, with emphasis on several individuals with CYTHA higher than the most productive parent. Broad-sense heritabilities ranged from intermediate (55.01% for FYTHA) to high values (80.50% for CI). Correlations between the predicted means were estimated (Figure 1). Low correlations (from 0.18** to 0.22***) were observed between FYTHA and root yield traits. The highest correlation (0.99***) was between CYTHA and RYTHA, which was expected, since most RYTHA is derived from CYTHA. Among the traits used for CI calculation, CYTHA also had the highest correlation with CI (0.79***), likely because it is its main component. TNR components were also highly correlated with TNR, namely NOCR (0.90***) and NONC (0.86***). Finally, NOCR and NONC turned out to be highly correlated with CYTHA (0.80***) and NCYTHA (0.84***), respectively.

**Table 1.**
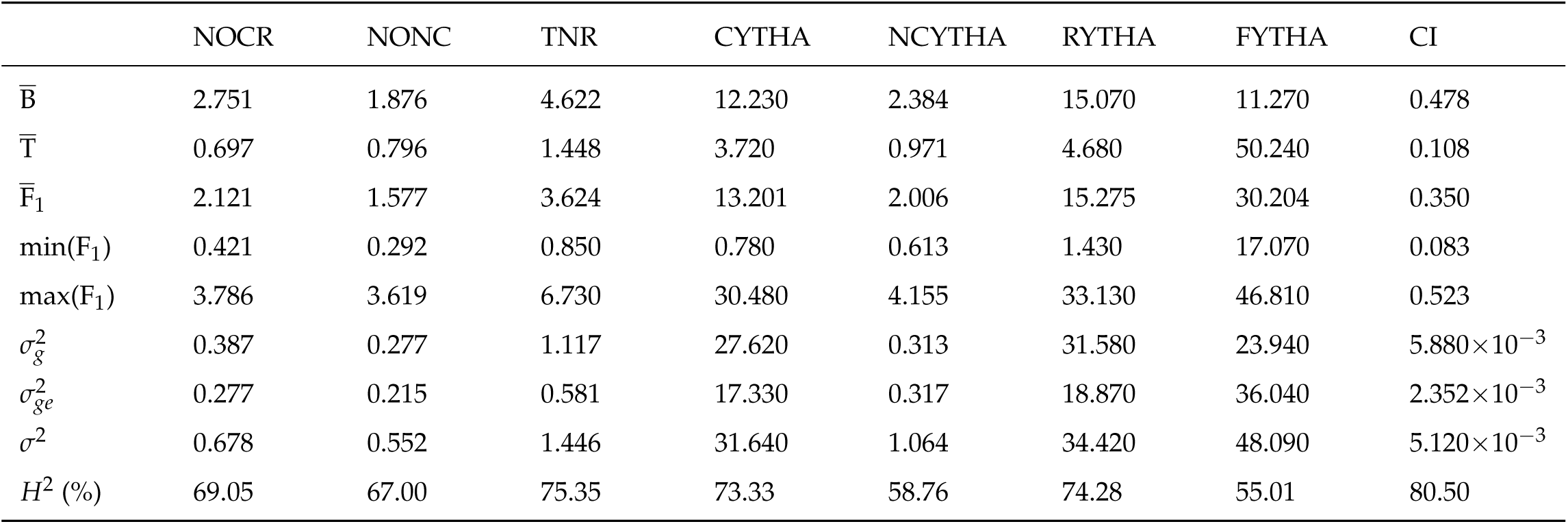
Phenotypic analysis summary of eight yield-related traits from ‘Beauregard’ × ‘Tanzania’ (BT) full-sib family. Parental (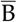 and 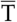) and progeny 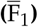 means, minimum and maximum F_1_ means, and genetic 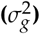, genotype-by-environment interaction 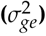 and residual (*σ*^2^) variance components and heritability (*H*^2^) estimates are shown for eight traits: number of commercial (NOCR), noncommercial (NONC) and total (TNR) roots per plant, commercial (CYTHA), noncommercial (NCYTHA) and total (RYTHA) root yield in t · ha^−1^, foliage yield (FYTHA) in t · ha^−1^, and commercial index (CI).

**Figure 1.**
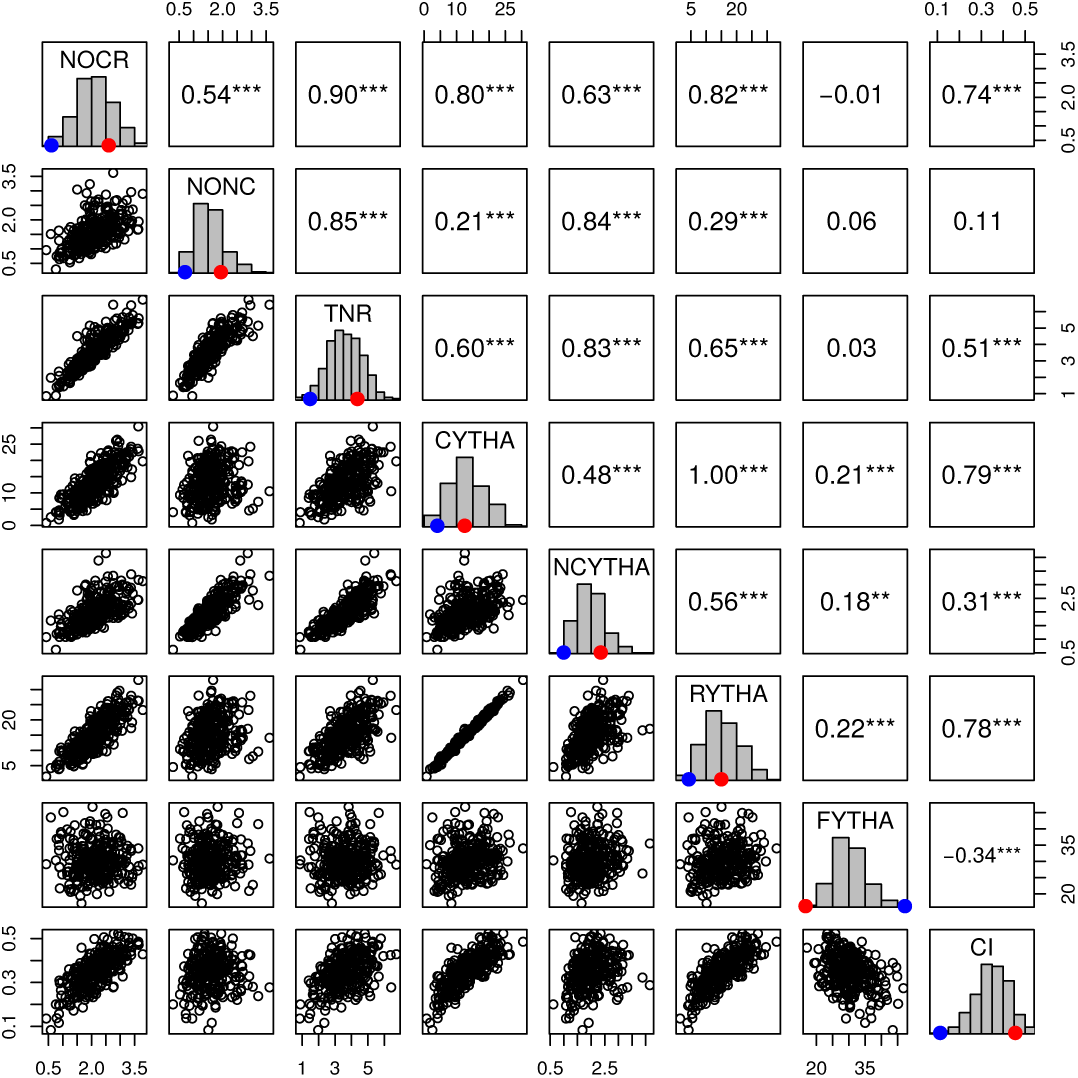
Pearson’s correlations (\*\**p* < 0.01, \*\*\**p* < 0.001) among predicted means of eight yield-related traits from ‘Beauregard’ × ‘Tanzania’ (BT) full-sib family. Parental means are represented by red (B) and blue (T) dots. Trait abbreviations: number of commercial (NOCR), noncommercial (NONC) and total (TNR) roots per plant, commercial (CYTHA), noncommercial (NCYTHA) and total (RYTHA) root yield in t · ha^−1^, foliage yield (FYTHA) in t · ha^−1^, and commercial index (CI).

### Mapping QTL in BT population

#### Simulations

The BT linkage map based on 298 F_1_ progenies was used to simulate 1,000 quantitative traits with three QTL each. We ran FEIM (Model 1) and REMIM (Model 3) for each simulated trait in order to assess their detection power and FDR in such scenario (see Tables S1 and S2). For REMIM, different forward *p*-value thresholds did not impact power or FDR (results not shown), but backward thresholds were critical. Figure 2 compares different threshold criteria for declaring a QTL during FEIM and REMIM (for *p* < 0.01 forward threshold). From both approaches, FDR greater than 20% were found when using less conservative criteria of genome-wide significance LOD threshold of 0.20 for FEIM and *p* < 10^−3^ backward threshold for REMIM. However, detection power was higher with REMIM (75.2%, on average) than with FEIM (65.6%, on average). Taking more conservative criteria such as genome-wide significance LOD threshold of 0.05 for FEIM and *p* < 10^−4^ backward threshold for REMIM, FDR decreased to 14.3% and 13.0%, respectively, whereas power was still higher with REMIM (67.3%, on average) than with FEIM (59.9%, on average). In fact, although the QTL with the highest heritability 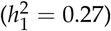 was similarly detected regardless of the method and criterion, a higher proportion of QTL with intermediate and low heritabilities were mapped under the multiple-QTL mapping approach. Interestingly, even with the most stringent criteria of *p* < 10^−5^ backward threshold for REMIM, we were able to map as many QTL as using genome-wide significance LOD threshold of 0.05 for FEIM, but with a better FDR control (∼10%). In general, the average absolute difference between the simulated and mapped QTL peak location did not differ when comparing models or thresholds whatsoever (see Table S1). From testing different *d* values for *LOD* − *d* and *LOP* − *d*, we learned that *d* = 1.5 was a good approximation of 95% support interval for both FEIM and REMIM (see Table S2).

**Figure 2.**
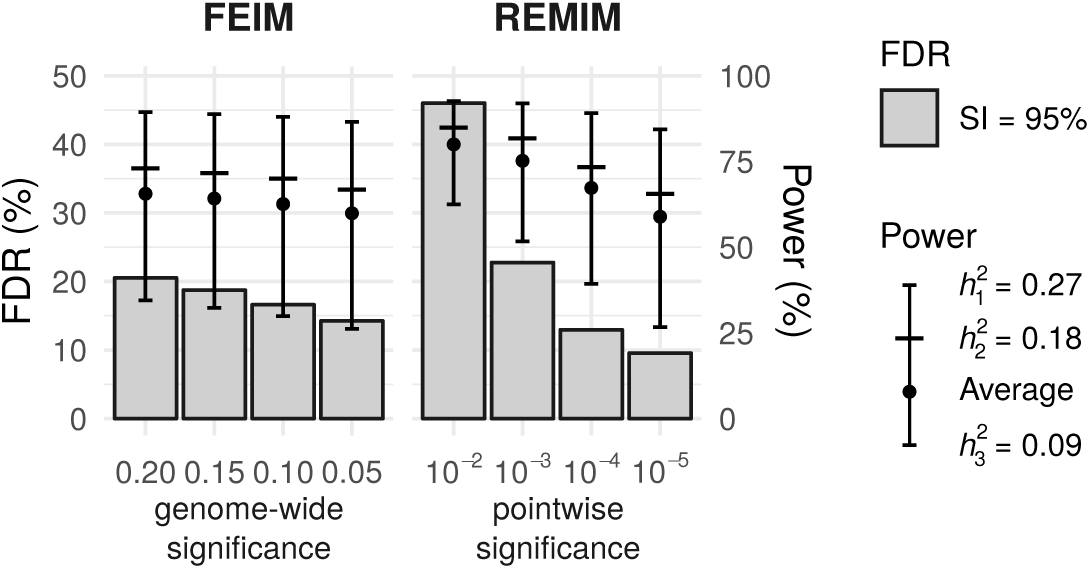
Detection power and false discovery rate (FDR) from QTL mapping analyses of 1,000 simulated traits in ‘Beauregard’ × ‘Tanzania’ (BT) full-sib family. Each trait was simulated with three QTL with different heritabilities 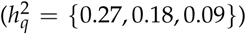, randomly positioned on the BT linkage map (*n* = 298). Fixed-effect interval mapping (FEIM) used different genome-wide significance LOD thresholds (0.20, 0.15, 0.10, 0.05) based on 1,000 permutation tests. Random-effect multiple interval mapping (REMIM) was performed using different score-based *p*-value thresholds during backward elimination (*p* < 10^−2^, 10^−3^, 10^−4^, 10^−5^) after forward search using *p* < 0.01. Power (vertical lines, right axis) represents the proportion of true QTL over the total number of simulated QTL. FDR (bars, left axis) depicts the proportion of false QTL over the total number of mapped QTL for a ∼95% support interval (SI) coverage.

#### Yield-related traits

We adopted 0.01 and 10^−3^ as the respective forward and backward *p*-value thresholds for detecting QTL in eight yield-related traits in the BT population using REMIM (Model 3; Figure 3, Table 2). In total, 41 QTL were identified, with *p*-values ranging from 1.64 × 10^−9^ (QTL 2 for TNR) to 7.20 × 10^−4^ (QTL 1 for RYTHA). The number of QTL per trait ranged from one (CYTHA and CI each) to ten (TNR). NOCR, NONC, NCYTHA, RYTHA and FYTHA had seven, nine, five, three and five QTL, respectively. All LGs except LG 5 harboured QTL regions. LGs 4 and 15 harboured six QTL each, and LGs 10 and 13 harboured four QTL each. Approximate 95% support intervals computed as *LOP* − 1.5 (see Figure S1) showed that QTL hotspots may be found on LGs 1, 3, 4 and 15 as several QTL regions seemed to be co-localized. On LG 1, QTL peaks for NOCR, TNR and NCYTHA can be found between 131.50 and 140.03 cM. On LG 3, QTL peaks for NONC, NOCR and TNR were localized either at 12.36 or 20.18 cM. LG 4 seemed to have a second QTL hotspot: a first region with QTL peaks between 32.06 and 76.48 cM for NONC, NOCR, TNR and FYTHA, in addition to a second region with another QTL for TNR and one for NCYTHA at 200.09 and 180.36 cM, respectively. Similarly, on LG 15, a first hotspot had QTL peaks either at 4.19 or 5.27 cM for CYTHA, RYTHA and CI, whereas a second hotspot had QTL peaks at 109.10 and 105.02 cM for TNR and NCYTHA, respectively.

**Table 2.**
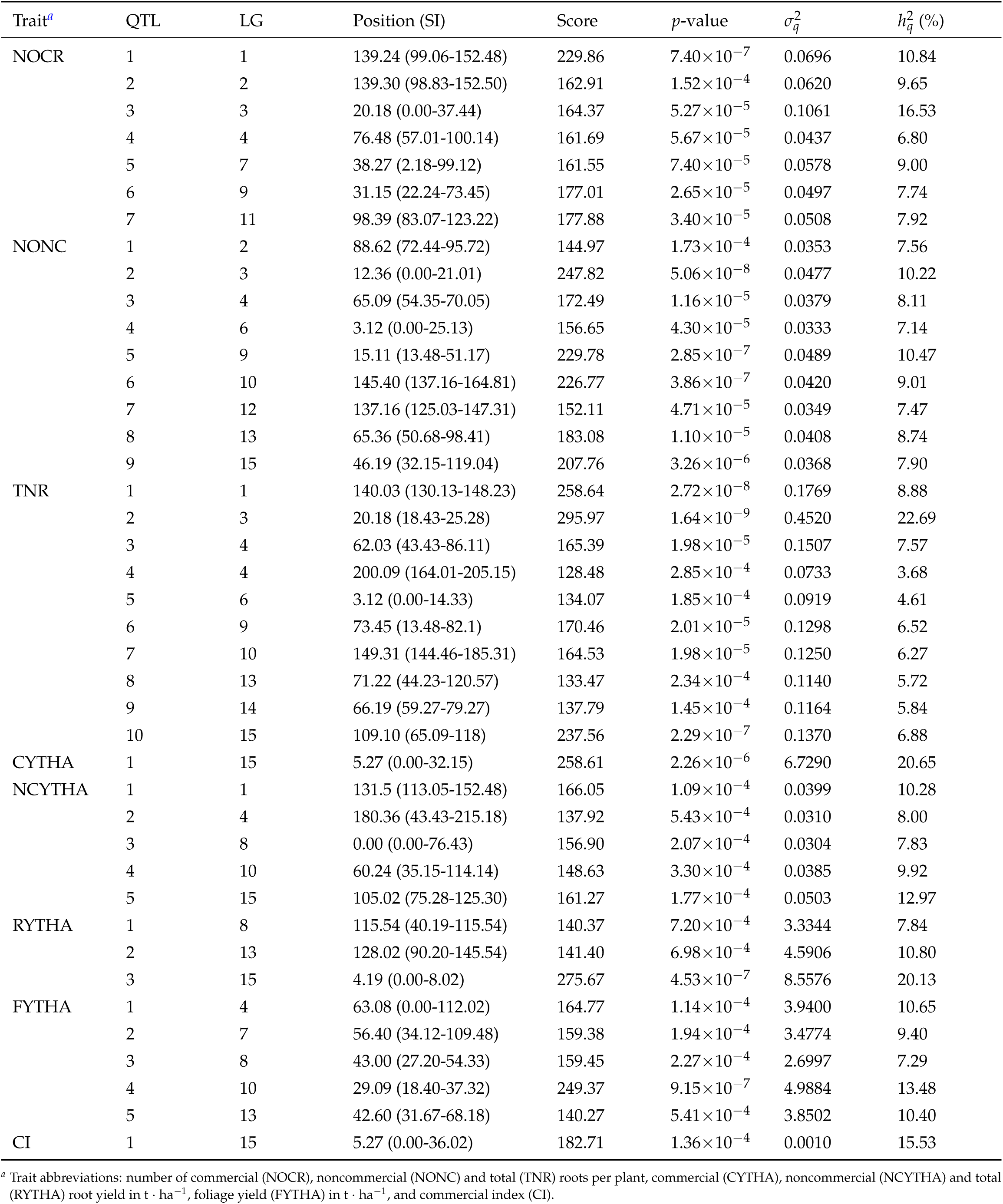
Random-effect multiple interval mapping (REMIM) of yield-related traits from ‘Beauregard’ × ‘Tanzania’ (BT) full-sib family. Linkage group (LG), map position (in centiMorgans) and its ∼95% support interval (SI), score statistic and its corresponding *p*-value, variance 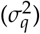 and heritability (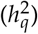, in percentage) of mapped QTL.

**Figure 3.**
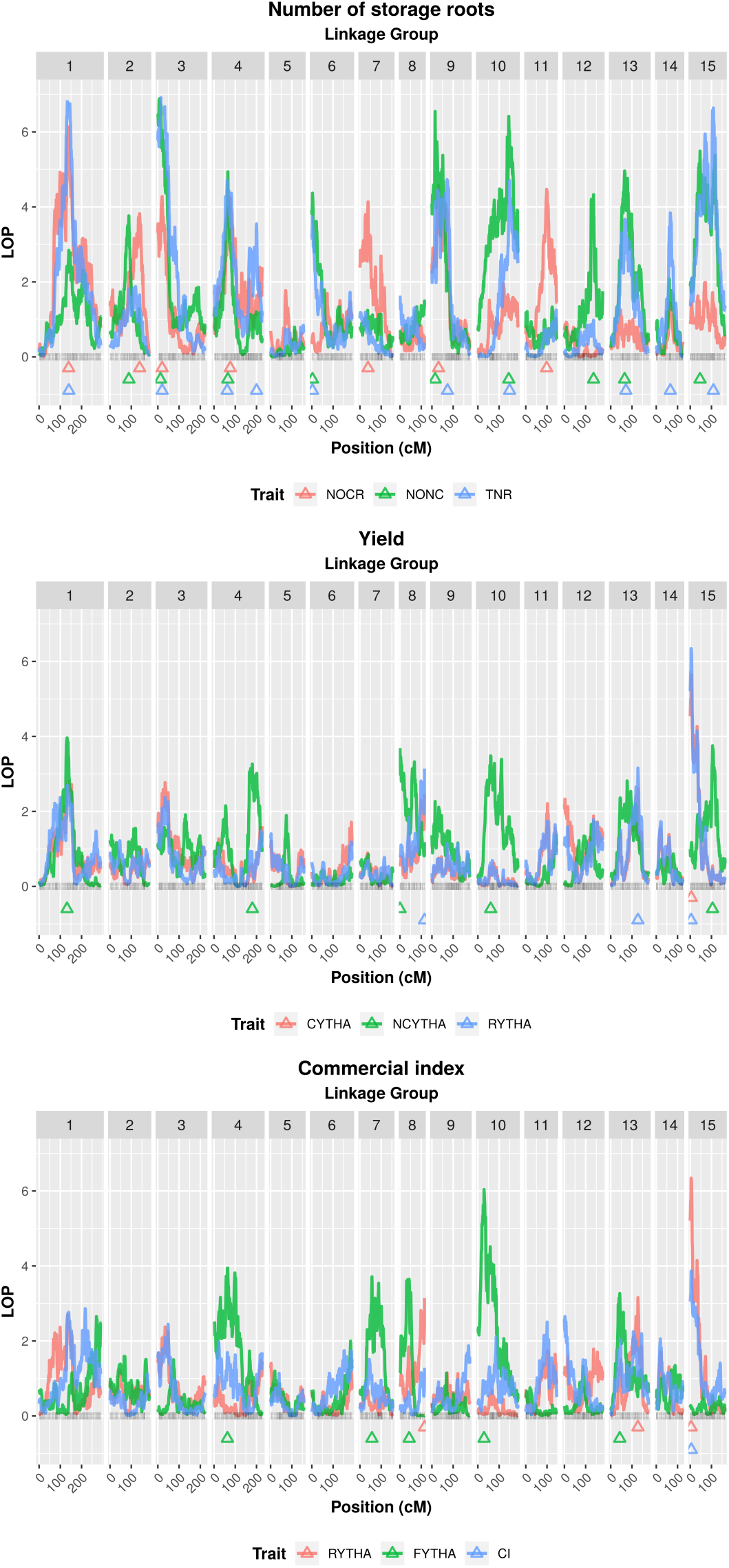
Logarithm of *p*-value (*LOP*) profiles from randomeffect multiple interval mapping (REMIM) of eight yield-related traits from ‘Beauregard’× ‘Tanzania’ (BT) full-sib family. Triangles show the QTL peak location. Trait abbreviations: number of commercial (NOCR), noncommercial (NONC) and total (TNR) roots per plant, commercial (CYTHA), noncommercial (NCYTHA) and total (RYTHA) root yield in t ha^-1^, foliage yield (FYTHA) in t ha^-1^, and commercial index (CI).

QTL variances 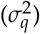 and heritabilities 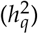 estimates from Model 2 are shown in Table 2, where the subscript *q* denotes the QTL number for a specific trait. QTL heritabilities ranged from 3.68 (QTL 4 for TNR) to 22.69% (QTL 2 for TNR), representing the proportion of the total variance explained by that QTL, conditional to all the other QTL in the model. Out of 41 QTL, 14 were considered major QTL 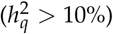, with one major QTL for TNR 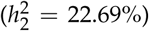, CYTHA 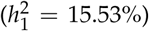, and CI 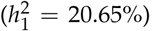, two major QTL for NONC (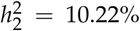 and 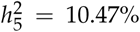), NOCR (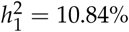 and 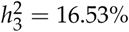), NCYTHA (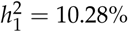 and 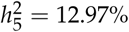), and RYTHA (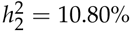 and 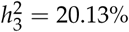), and three major QTL for FYTHA (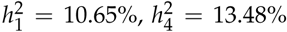 and 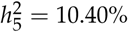). Altogether, mapped QTL explained as much as 76.64%, 68.49% and 78.66% of the total variance for NONC, NOCR and TNR, respectively. As less QTL were identified for root yield traits, a relatively smaller portion of the total variance was explained for NCYTHA (48.99%), RYTHA (38.77%) and FYTHA (51.22%). Interestingly, most of the major QTL lies on hotspots, such as those on the beginning of LGs 3 and 15 (see Figure S2). In order to compare QTL detection results, we used a rather relaxed genome-wide significance of 0.20 for FEIM, whose permutation-based LOD score thresholds ranged from 6.81 to 6.89 depending on the trait (*LOD* ≈ 6.85, on average) (see Figure S4). A total of 17 QTL were mapped (see Table S3): one for each CYTHA, RYTHA and CI, two for each NOCR, NCYTHA and FYTHA, and four for each NONC and TNR. Six LGs harboured QTL: LGs 15 and 3 had six and four QTL, respectively, LG 1 had three QTL, LG 10 had two QTL each, and LGs 4 and 9 had only one QTL each. In fact, the most significant QTL (*LOD >* 9) were found on LGs 1, 3 and 15, similar to REMIM results. In addition to QTL not detected on several other LGs, FEIM did not detect any QTL for number of roots on LG 4, identified as a QTL hotspot by REMIM, although for FYTHA a QTL reached the threshold at 95.01 cM (*LOD* = 6.91). On the other hand, REMIM missed a QTL for NCYTHA on LG 3 at 32.56 cM, which was detected using FEIM. LOD thresholds for a genome-wide significance of 0.05 ranged from 7.68 to 7.98 (*LOD* ≈ 7.77, on average), and one would have mapped 11 QTL, instead. One QTL of each NONC (LG 1), TNR (LG 9), NCYTHA (LG 3) and RYTHA (LG 15), and both QTL for FYTHA (LGs 4 and 10) would have been missed under this more conservative criterion.

From REMIM, allele and allele combination additive effects of each QTL (see Table S4) were derived from the QTL genotype BLUPs for each trait (Model 2). These effects represent the parental contribution to the population mean, i.e. how much one adds to or subtracts from the mean given one of the 400 possible genotypes. For instance, Figure 4 shows the allele-specific and allele combination additive effects of QTL 1 for CYTHA. Inferences on which alleles contribute more to the mean as well as which ones ought to be selected for breeding purposes are straightforward. For example, individuals with the haplotypes *b* from ‘Beauregard’ and *i* from ‘Tanzania’, and without the haplotypes *c* and *j* through *l* from the respective parents will have the highest QTL-based breeding value estimates for CYTHA. In order to allow comparison among the allele effects from 41 QTL, proportional contribution of a specific allele was calculated as the ratio between its absolute effect and the sum of all 12 absolute effects for each QTL, so that effects would range within the unit interval. Out of 492 effects, 75 (15.24%) showed a contribution of at least 15% (almost double of the average). Among these most important effects, ‘Beauregard’ provided 34 effects, with 15 negative and 19 positive effects summing up to -2.833 and +3.908, while ‘Tanzania’ contributed with 41 effects, with 22 negative and 19 positive effects summing up to -4.308 and +3.594. Although most QTL had approximately 50%:50% allele effect contribution from each respective parent, we observed skewed contributions towards ‘Beauregard’ (from 61%:39% to 72%:28%) for seven QTL, all related with number of roots, and towards ‘Tanzania’ (from 40%:60% to 30%:70%) for eight QTL, with four related to number of roots and four to yield traits. By computing QTL-based breeding values we could hypothesize on the genetic basis of trait correlation. For example, the single locus (QTL 1) detected for CI has shown additive effects in the same direction as those of QTL 1 for CYTHA. Therefore, correlation between QTL-based breeding values from these two traits were indeed as high as 0.97*** (see Figure S3). In addition, other important breeding lessons may be learned from these correlations. For instance, very low, non-significant correlation between CYTHA and NCYTHA (0.11) indicates that improving the former will not affect the latter. An increase of NOCR would bring a relative increase of NONC, though, as their QTL-based breeding values showed higher, significant correlation (0.46***). More interestingly, QTL-based breeding values for FYTHA did not seem to correlate to any other root-related trait. Finally, the absolute positive correlation of 1.00*** between predicted means from CYTHA and RYTHA (Figure 1) could be only partially explained by a single co-localized QTL, since the correlation between QTL-based breeding values was smaller (0.73***), although still high.

**Figure 4.**
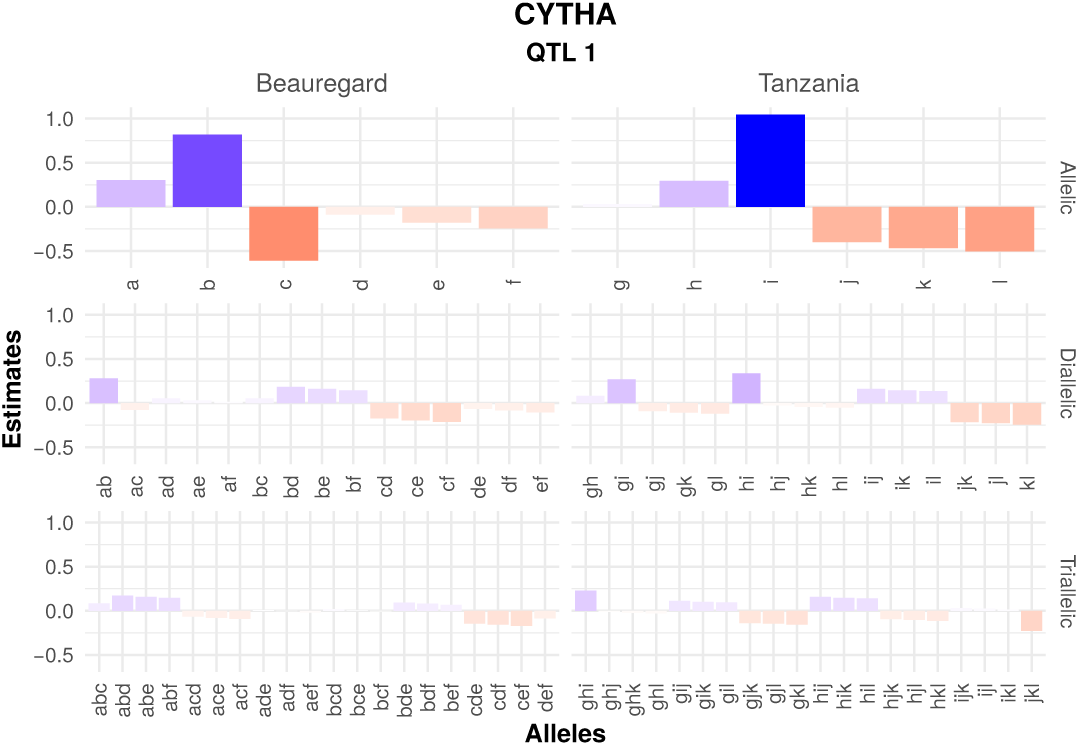
Allele and allele combination additive effects from the decomposed best linear unbiased predictions (BLUPs) for the QTL 1 (on linkage group 15 at 5.27 cM) of commercial root yield in t · ha^-1^ (CYTHA) in a hexaploid sweetpotato full-sib family (‘Beauregard’× ‘Tanzania’). Marker-assisted selection for increasing CYTHA would have to focus on selection for alleles *b* and *i*, and against alleles *c* and *j* through *l* from the respective parents.

#### Candidate genes underlying QTL hotspots

This study identified numerous QTL corresponding to yield-related traits and identification of causal genes within all QTL will require further research efforts due to the large number of genes that are involved in storage root formation and yield underlying these QTL. As we have access to an expression profiling dataset for ‘Beauregard’ and ‘Tanzania’ roots, we elected to examine putative candidate genes under two QTL that had high heritability and effect sizes: the QTL for TNR on LG 3 (colocalized for NOCR and NONC) and the QTL for CYTHA on LG 15 (co-localized for RYTHA and CI).

The QTL peak for TNR on LG 3 was at 1,591,872 bp with a SI between 1,428,660 and 1,971,958 bp (see Table S5) that contains 75 genes. Examination of functional annotation of these 75 genes, coupled with expression profiles in leaves as well as a time course of developing roots in both ‘Beauregard’ and ‘Tanzania’ (Gemenet *et al*. 2019, submitted) revealed three candidate genes of interest (see Figure S5). The *I*. *trifida itf03g02930* gene encodes a homolog of SKU5, a glycosyl phosphatidylinositol modified protein in *Arabidopsis thaliana* with similarity to multiple-copper oxidases that are localized to the plasma membrane and cell wall. In sweetpotato, the *itf03g02930* homolog was expressed in leaves and roots, although the expression in roots is substantially higher than that in leaves. A second candidate gene is *itf03g03280* which encodes a protein with sequence similarity to annexin. The *itf03g03280* homolog shares 71% identity (85% similarity) with 99% coverage with ANN2 (AT5G65020) and 66% identity (81% similarity) over 100% coverage with ANN1 (AT1G35720). *itf03g03280* was expressed in both leaves and roots (total roots, fibrous roots, and storage roots) although expression in roots is nearly twice that of leaves. The last candidate gene, *itf03g03460*, encodes a protein with 51% identity (62% similarity) to the WUSCHEL homeobox family protein (AtWOX13). While *itf03g03460* was lowly expressed in leaves, it was highly expressed in roots of both ‘Beauregard’ and ‘Tanzania’.

On LG 15, a major QTL for CYTHA with the peak at 477,772 bp spanned positions from 21,822 to 1,939,509 bp (see Table S5) and 310 genes. As this was too large of a distance to manually curate candidate genes responsible for the trait, we restricted our query to 25 genes distal and proximal to the most significant marker. Within this region, two genes encoded functions that may be associated with storage root development and had expression profiles that supported a role in storage root development (see Figure S5). The hormone ethylene has diverse roles in cell proliferation and elongation, and the *I*. *trifida itf15g01020* gene encodes a protein with similarity to the *A*. *thaliana CONSTITUTIVE TRIPLE RESPONSE 1* gene (*CTR1*) with functions in the ethylene signaling pathway. In sweetpotato, the *itf15g01020* homolog was expressed in leaves but expressed at twice the levels in developing roots. Storage roots are grown for their high starch content, and *itf15g01120* encodes a protein with similarity to starch branching enzyme 2.2, involved in starch biosynthesis. *itf15g01120* was expressed in leaves and roots with the highest expression levels detected in storage, not fibrous or developing roots.

## Discussion

Most of the linkage and QTL mapping work done for sweet-potato so far has relied on strategies based on a double pseudo-testcross approach for diploid species (Grattapaglia and Sederoff 1994). For example, separate parental maps have been built based on this diploid-based simplification, using qualitative marker systems such as randomly amplified polymorphic DNA (RAPD; Ukoskit and Thompson 1997), amplified fragment length polymorphism (AFLP; Kriegner *et al*. 2003; Cervantes-Flores *et al*. 2008a; Nakayama *et al*. 2012), retrotransposon insertion polymorphisms (Monden *et al*. 2015) and simple sequence repeats (SSR; Kim *et al*. 2017). A recent map was developed from a selfing population and used only single-dose SNPs, resulting in higher marker saturation in comparison to the previous maps (Shirasawa *et al*. 2017), though the map was still not integrated. In some of these cases, QTL mapping analyses were performed for several traits, mostly related to quality (Cervantes-Flores *et al*. 2011; Zhao *et al*. 2013; Yu *et al*. 2014; Kim *et al*. 2017) and resistance to biotic stresses (Cervantes-Flores *et al*. 2008b; Yada *et al*. 2017a). For yield-related traits, only two studies have been reported to date (Chang *et al*. 2009; Li *et al*. 2014). The use of DNA markers with unknown DNA sequence limited our ability to compare their results with *I*. *trifida* and *I*. *triloba* genomes (Wu *et al*. 2018), and ultimately with our present QTL study (see Table S5). Moreover, although these diploid-based strategies were the state-of-the-art at that time for qualitative marker-based, low density genetic maps, they imposed significant restrictions on statistical power for QTL detection and its genetic interpretation. Recently, more improved methods and computational tools that take into account autopolyploid complexity for dosage SNP calling (Voorrips *et al*. 2011; Serang *et al*. 2012; Schmitz Carley *et al*. 2017; Gerard *et al*. 2018) and integrated linkage map construction (Hackett *et al*. 2016; Bourke *et al*. 2018; Mollinari and Garcia 2018) have become available, mostly dedicated to tetraploids. Taking advantage of the newly developed MAPPOLY package, (Mollinari *et al*. 2019, in preparation) built the first integrated genetic map for sweetpotato, from the BT population used here. For a hexaploid species, this has opened up new opportunities for more interpretable QTL genetic models due to MAPPOLY implementation of a HMM that delivers QTL genotype conditional probabilities along a fully integrated genetic map (Mollinari and Garcia 2018).

More specifically, QTL mapping in autopolyploid species has been limited to a fixed-effect interval mapping (FEIM) model proposed for tetraploids (Hackett *et al*. 2001) and eventually expanded for hexaploids (van Geest *et al*. 2017). Consisting of a single-QTL model, 2*m*-2 main effects are fitted (*m* is the ploidy level), and this model is compared to a null model (with no QTL) using LRT, ultimately expressed as LOD scores. Permutation-based genome-wide significance LOD thresholds are then used to declare a QTL. Trying to add more QTL into FEIM could rapidly lead the model to over-parameterization, since each QTL requires as much as six (for tetraploids), ten (for hexaploids) or 14 (for octoploids) parameters to be estimated. Furthermore, new rounds of permutation, based on a model with QTL, would need to be carried out in order to provide an updated LOD score threshold (Klaassen *et al*. 2019). In contrast, the random-effect multiple interval mapping (REMIM) model presented here is designed to fit multiple random-effect QTL by estimating only one single parameter 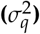 per QTL. Score statistic tests are performed in order to assess whether a QTL variance component is zero or not, conditional to other QTL in the model. These tests provide an approach for comparing two nested models with the reduced model having a random effect excluded, just like residual LRT (RLRT) would do. However, (R)LRT is more prone to numerical errors because the null hypothesis 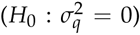 falls on the boundary of the parameter space, whereas score-based methods can be robust to eventual misspecification of the distribution of random effects (Verbeke and Molenberghs 2003).

We used the BT population genetic map to simulate quantitative traits in order to compare FEIM and REMIM performances, and also to assess the impact of using different thresholds for QTL detection (Figure 2). According to our simulations, both approaches would detect similar number of simulated QTL at more stringent criteria, with REMIM delivering less false positives. The results also suggested that one could use a more relaxed criteria in order to increase the power of detection while still maintaining an acceptable level of FDR. Although conclusions may be limited to the simulated scenario, multiple-QTL model approaches have been proven to provide greater power and better FDR control than single-QTL models for both univariate (Zeng *et al*. 1999; Laurie *et al*. 2014) and multivariate models (Da Costa E Silva *et al*. 2012b), mostly due to the differences in detecting QTL with smaller effects. In fact, this is rather expected as a multiple-QTL model has a smaller residual variance which helps to detect additional QTL. Multiple-QTL models are also supposed to improve detection of more than one QTL on the same LG (Mayer 2005), as they are usually hard to separate from each other due to the high extension of linkage disequilibrium in mapping populations. For polyploids, a non-optimal approach of using residuals from a fitted single-QTL model as phenotypic data to find a second linked QTL has been proposed (Mengist *et al*. 2018). In QTL mapping analysis, it is important to have a reasonable balance between detection power and FDR, as we are interested in mapping as many true QTL as possible. It should also consider the goals of the study, i.e. whether it is intended to use a few very reliable QTL for marker-assisted breeding, or to discover as many QTL-related putative genes as possible for further validation. Additional suggestive QTL also increase the number of hypothesized regions that affect trait variation and may be targeted for selection. For yield-related traits in our BT population, FEIM results were limited to 15 QTL (see Figure S4), most of which also happened to be mapped using REMIM (Figure 3). In fact, by performing REMIM, we found 27 minor QTL 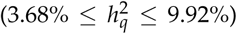 in addition to 14 major ones 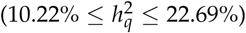 (Table 2, see Figure S2). Based on double pseudo-testcross approach, previous estimates of proportion of variance explained (PVE) by nine QTL for storage root yield ranged from 17.7 to 59.3% for 202 individuals from a cross between two Chinese sweetpotato varieties (Li *et al*. 2014). In another study, analyses of two reciprocal full-sib populations with less than 120 individuals each detected seven QTL for root and top (foliage) weight (16.0% ≤PVE ≤29.5%), and only one QTL detected for root number (PVE = 14.8%) (Chang *et al*. 2009). Because of likely estimation bias due to reduced sample size and the use of not very informative markers and linkage maps, these previous PVE findings are hard to compare with our results. Adjusted *R*^2^ from FEIM accounted for 7.8% ≤ PVE ≤12.6% (see Table 3), which may not be comparable with *h*^2^ from REMIM due to different approaches (single vs. multiple QTL models).

In general, the number of mapped QTL was compatible with the proportion of total variance explained by the QTL 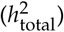 (Table 2). From seven to ten QTL 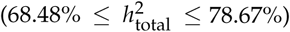 were mapped for traits related with number of roots (67.00% ≤ *H*^2^ ≤ 75.35%), whereas from one to five 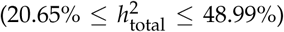 were found underlying root yield variation (58.76% ≤ *H*^2^≤ 74.28%). Finally, there were five QTL 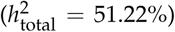 for FYTHA (*H*^2^ = 55.01), but only one 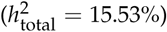 for CI (*H* = 80.50). Although number of roots seemed to be as heritable as root yield (Table 1), the latter traits are likely more complex in terms of their genetic architecture than the former ones. That is, not only number of roots contributes to yield, but also size and composition, so we can expect that more regions are involved in root yield, in addition to those involved in number of roots. Nevertheless, Yada *et al*. (2017b) found a rather low trait heritability (likely individual-basis) for commercial root yield (*H*^2^ = 24%) among 278 full-sibs of a cross between ‘New Kawogo’, a Ugandan landrace, and ‘Beauregard’, possibly due to stronger G×E interaction, which adds to the trait complexity. Here, G×E interaction seemed important for all traits and its consequences to QTL mapping and breeding will be explored in future studies. As QTL mapping targets major QTL, usually stable across environments, most of the minor ones must have gone undetected. Moreover, additional genetic variation could be due to higher order allele interactions and genetic epistasis, which the current models do not account for. In fact, only a few minor QTL co-localized among number of roots and yield traits, which explains lower correlations among QTL-based breeding values (from -0.02 to 0.63***, see Figure S3) relative to correlations among predicted means (from 0.21*** to 0.84***, Figure 1) between these sets of traits. Based on the correlation between QTL-based breeding values, FYTHA does not seem to be useful in indirect selection for CYTHA (−0.02, see Figure S3), as suggested previously (Chang *et al*. 2009), even though some correlation (0.21**) among their predicted means was observed (Figure 1). Although ‘Beauregard’ and ‘Tanzania’ contributed more importantly with positive and negative major effects, respectively, the presence of both favorable and unfavorable QTL alleles in either parents possibly explains the presence of transgressive segregants for all traits. Transgression in polyploids seems to be due to not only cumulative complementary alleles at different loci (Tanksley 1993), but also from the same QTL. In fact, increased heterozygosity has been suggested as one of the major forces of polyploid evolutionary success, as a broader allele repertoire may result in the variation of gene expression and regulation needed to thrive in more diverse environmental conditions (Van De Peer *et al*. 2009). As an example, ‘Tanzania’ exhibited allele contributing to increase CYTHA from a major QTL (Figure 4), although this landrace was not very productive in our environments overall. The additive effects are the most important when performing selection as for a breeding point-of-view. However, one could easily estimate eventual dominance effects from the detected QTL using simpler biallelic-based models as proposed previously (Hackett *et al*. 2014; Chen *et al*. 2018). The effective use of higher allele interactions in QTL detection remains limited, though.

Several studies have looked at genes involved in storage root initiation and development in sweetpotato as reviewed by Khan *et al*. (2016). The storage roots differentiate from lateral roots by development of cambia around the protoxylem and econdary xylem, while lignification of the steles of some lateral oots inhibits this transformation (Villordon *et al*. 2012). The transformation is genetically and environmentally controlled. Using the expression profile of the parents of the current map ing population, we found genes in leaves and roots (see Figure S5) related to root development and sugar transport within the QTL hotspot on LG 3 associated with number of storage roots, implying that both root restructuring and carbon supply is likely involved in the number of lateral root that transform to storage root. In A. *thaliana*, SKU5 (*itf03g02930* homolog) was shown to have a role in regulating directional root growth as mutants in SKU5 were shorter than wild-type and altered in the angle of tip growth when grown on agar (Sedbrook *et al*. 2002). ANN1 and ANN2 (*itf03g03280* homolog) have been shown to be involved in post-phloem sugar transport to the root tip (Wang *et al*. 2018), a phenotype essential to development of storage roots. AtWOX13 (*itf03g03460* homolog) was shown to take part in lateral root development (Kreis *et al*. 2008). Other genes such as *SRF1* through *SRF10* (Tanaka *et al*. 2005), *knotted1*-like homeobox (*KNOXI*; Tanaka *et al*. 2008), MADS-box genes (Kim 2002), expansin (EXP) genes and BEL1-like homeodomain (Ponniah *et al*. 2017) have been strongly implicated in storage root formation and development in sweetpotato. Though we did not find evidence of differential expression of these genes in the available transcriptomic data, two MADS-box transcription factors (*itf03g02230* and *itf03g02240*), a BEL1-like homeodomain (*itf03g02670*) and an EXP (*itf03g05010*) were all found within the QTL region on LG 3. The association between these genes and the genes described in this study is yet to be defined and suggests the complex nature of storage root formation and development. On the QTL hotspot related to storage root weight on LG 15, we found the *CTR1* gene (*itf15g01020* homolog), which encodes a serine-threonine kinase and functions in the ethylene signaling pathway leading to inhibition of cell proliferation (Ramzan *et al*. 2015) had variable expression in the sampled roots. Rose *et al*. (1997) showed that inhibition of ethylene biosynthesis led to inhibition of *EXP1* gene in tomato. In sweetpotato, down-regulation of an *EXP1* homologue (*IbEXP1*) enhanced storage root development (Noh *et al*. 2013). While little is known about the role of ethylene in storage root development, the complex interactions of multiple hormones in storage root formation would suggest ethylene may be involved in storage root development. The main component of the sweetpotato storage root is starch. We found differential expression of a gene encoding starch branching enzyme (*itf15g01120* homolog). Starch biosynthesis involves four major classes of enzymes: ADP-glucose pyrophosphorylases, starch synthases, starch branching enzymes and starch debranching enzymes (Li *et al*. 2014). Starch branching enzymes influence the structure of starch through formation of *α* -1,6-branch points with different frequencies and chain length (Tetlow and Emes 2014). Given the number and magnitude of QTL identified in the current study as associated with yield and yield component traits, the results indicate that the candidate genes identified in the current study may interact with those from other loci to determine the final yield in terms of number, composition and weight of storage roots.

Here, we present a stepwise-based algorithm for multiple-QTL model selection in full-sib populations of autopolyploid species with a fully integrated map, from which QTL genotype conditional probabilities can be calculated. The use of score statistics is a key component of this new method, which depends on a dynamic and fast-computing test for model selection during the QTL search process. Simulations were performed in order to assess the impact of using different threshold criteria for QTL detection and to provide some empirical sense on how to use the method in practice. REMIM has been carried out in a hexaploid sweetpotato population to detect both minor and major loci contributing to the variation of yield-related traits that may be targeted in molecular-assisted breeding. The use of random-effect models has created the context for fitting multiple QTL, providing straightforward information on variance components, important for computing QTL heritabilities. Finally, QTL genotype predictions allowed us to estimate allele-specific additive effects, for characterizing major additive allele contributions, and compute QTL-based breeding values, that can be used for performing selection. This novel approach may enable more complex models, such as those accounting for interaction among QTL as well as multiple traits or multiple environments in order to study shared genetic control in different traits/environments and G ×E interaction at QTL level. Understanding the genetic architecture of root yield and other traits related to quality and resistance to biotic and abiotic stresses represents great opportunity for improving interesting characteristics in sweetpotato and other polyploids. Most of these important traits are polygenic in nature and only assessed later in a breeding program, where marker-assisted selection could help to speed up the process.

## Author contributions

DCG, WJG, AK, GCY and ZBZ conceived and designed the experiments. BAO performed DNA sequencing. DCG, FD, VM, WJG and AK carried out field experiments. GSP, MM and ZBZ developed tools and analyzed the data. CRB, JCW and DCG carried out candidate gene expression profiling. GSP wrote the manuscript. All authors read and approved the manuscript.

## Acknowledgments

This work was supported by the Bill & Melinda Gates Foundation [OPP1052983] as part of the Genomic Tools for Sweet-potato (GT4SP) Project. We acknowledge CIP sweetpotato breeding technical team in Peru for running experiments and collecting phenotypic data. Research at CIP was undertaken as part of the CGIAR Research Program on Roots, Tubers and Bananas (RTB), which is supported by CGIAR Fund Donors (http://www.cgiar.org/about-us/our-funders/).

## Supplemental Figures

**Figure S1.**
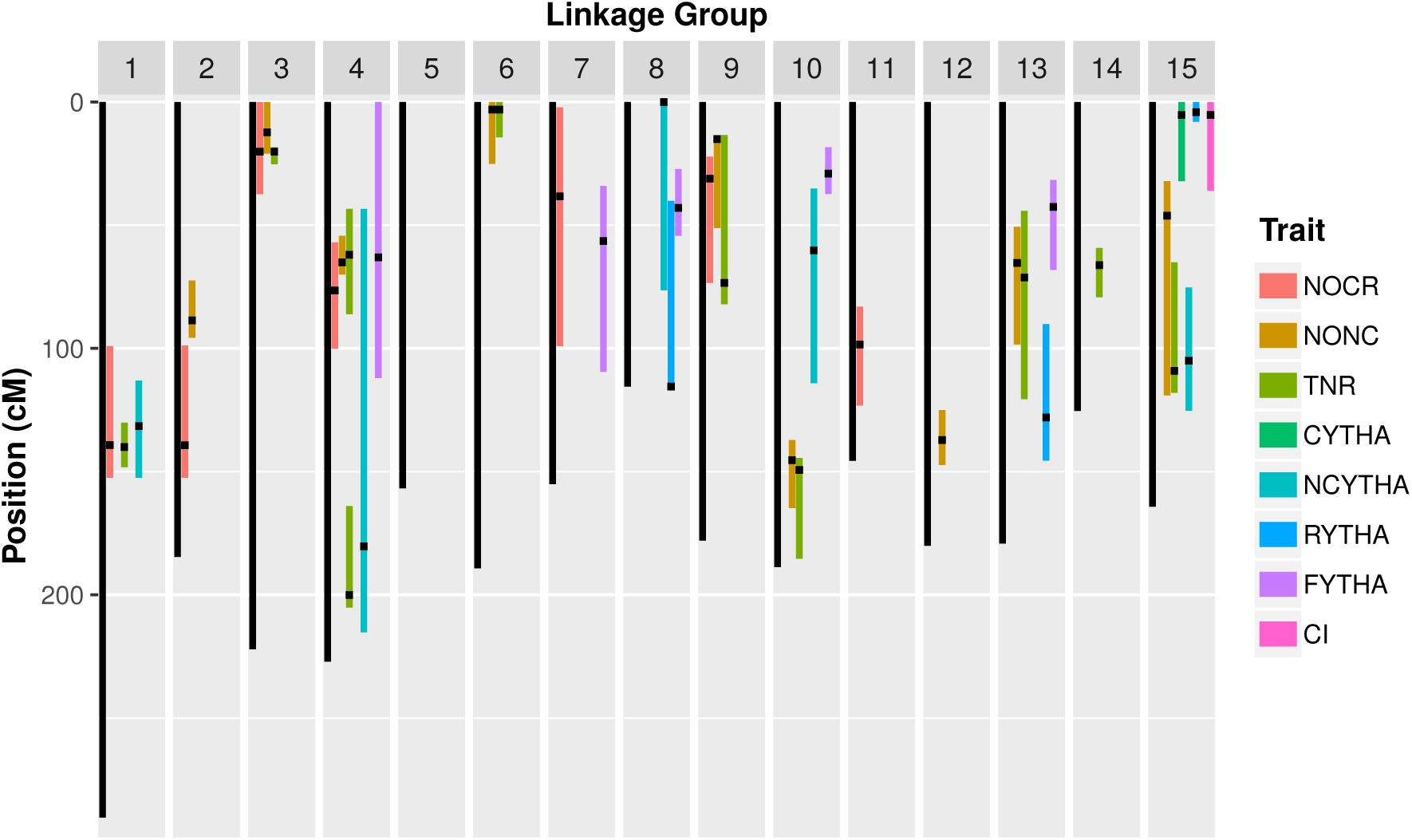
QTL support intervals from random-effect multiple interval mapping (REMIM) of yield-related traits from ‘Beauregard’ *×* ‘Tanzania’ (BT) full-sib family. Black dots represent the QTL peaks, and colored bars represent the ∼95% support interval computed as *LOP -* 1.5. Trait abbreviations: number of commercial (NOCR), noncommercial (NONC) and total (TNR) roots per plant, commercial (CYTHA), noncommercial (NCYTHA) and total (RYTHA) root yield in t *·* ha^*-*1^, foliage yield (FYTHA) in t *·* ha^*-*1^, and commercial index (CI).

**Figure S2.**
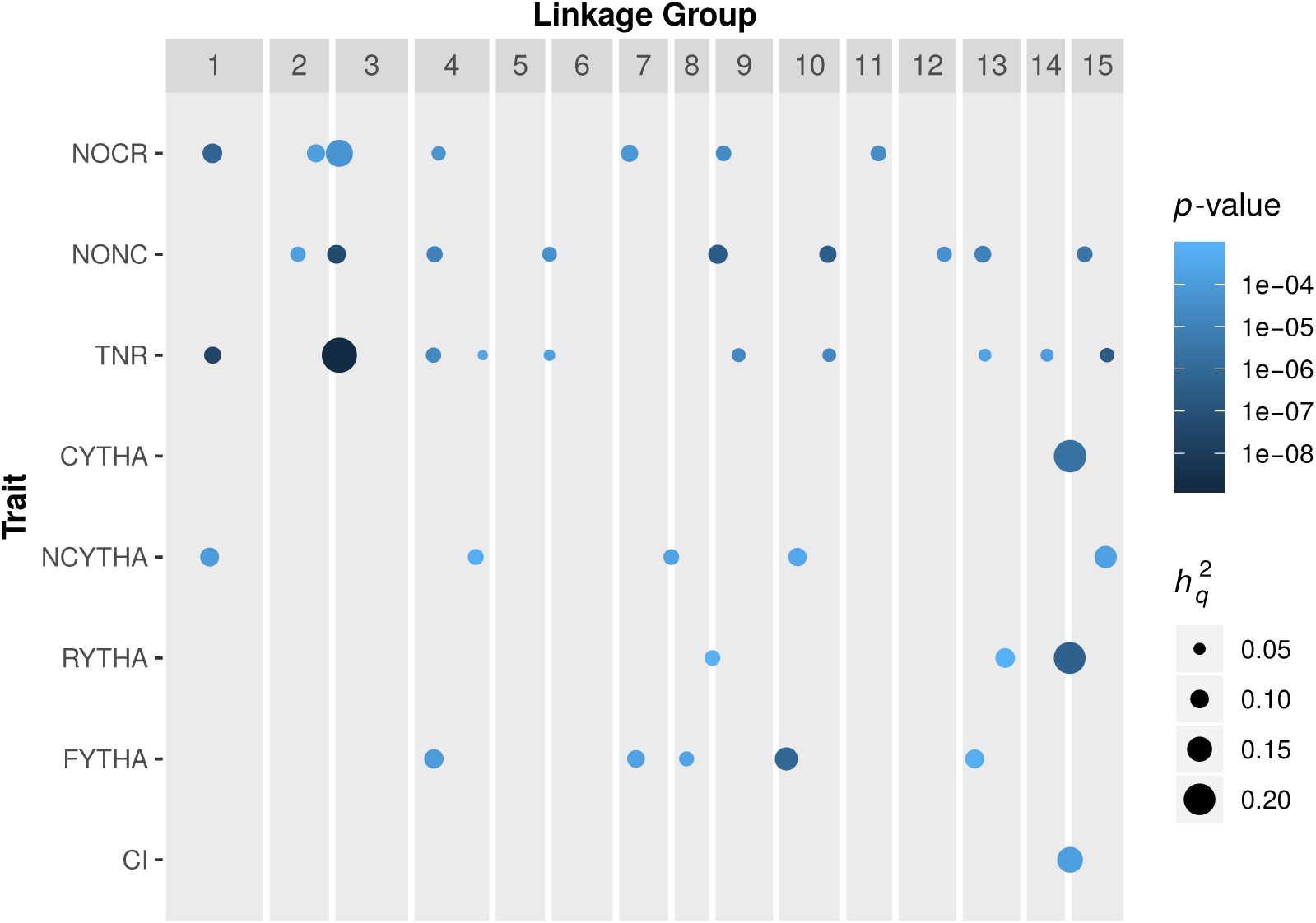
Score-based *p*-values and QTL heritabilities 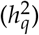 from random-effect multiple interval mapping (REMIM) of eight yield-related traits from ‘Beauregard’ *×* ‘Tanzania’ (BT) full-sib family. Dots are positioned relative to the QTL peaks: color gradient represents the *p*-values, while sizes are proportional to heritabilities of mapped QTL. Trait abbreviations: number of commercial (NOCR), noncommercial (NONC) and total (TNR) roots per plant, commercial (CYTHA), noncommercial (NCYTHA) and total (RYTHA) root yield in t *·* ha^*-*1^, foliage yield (FYTHA) in t *·* ha^*-*1^, and commercial index (CI).

**Figure S3.**
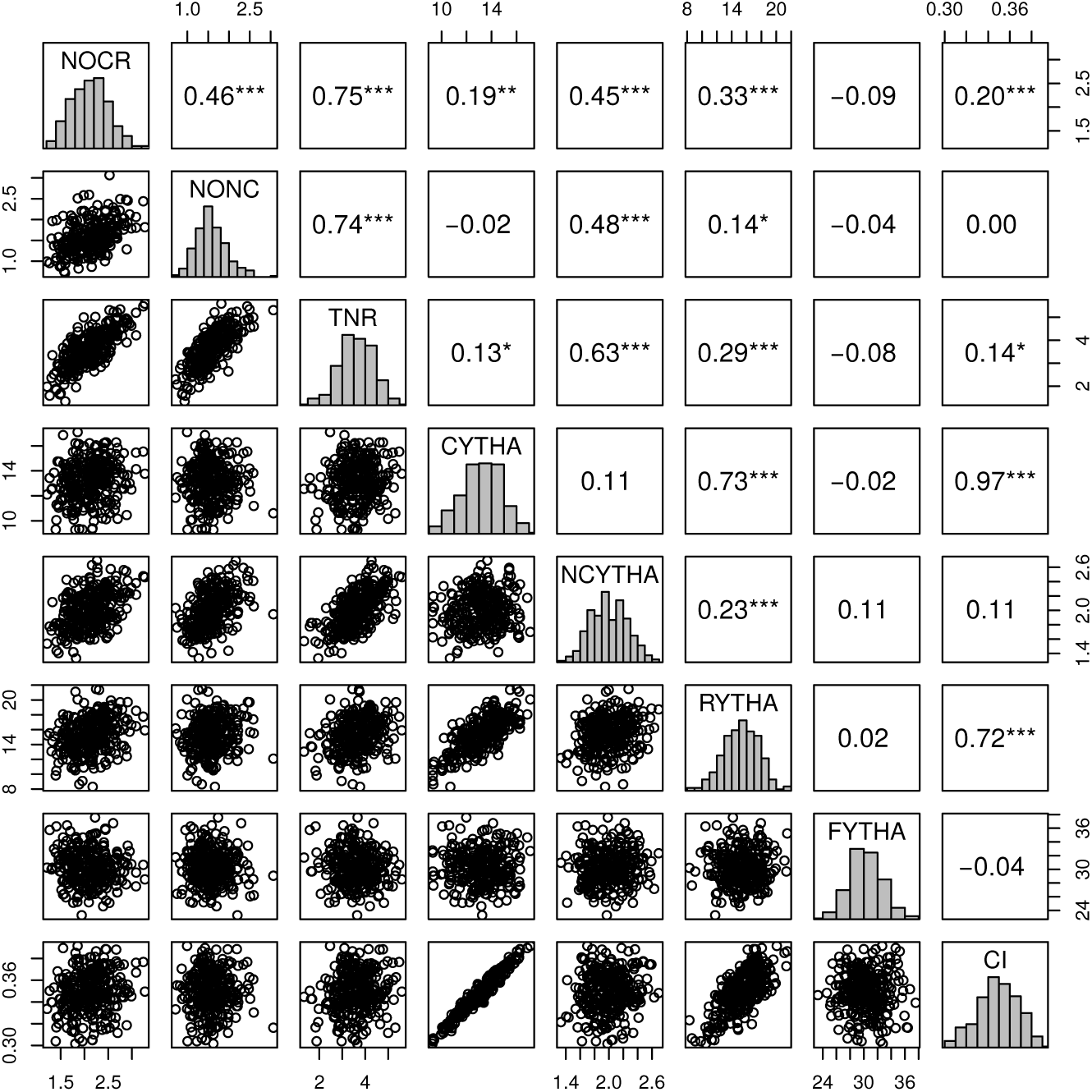
Pearson’s correlations (\**p <* 0.05, \*\**p <* 0.01, \*\*\**p <* 0.001) among QTL-based breeding values for eight yield-related traits from ‘Beauregard’ × ‘Tanzania’ (BT) full-sib family. Trait abbreviations: number of commercial (NOCR), noncommercial (NONC) and total (TNR) roots per plant, commercial (CYTHA), noncommercial (NCYTHA) and total (RYTHA) root yield in t *·* ha^*-*1^, foliage yield (FYTHA) in t *·* ha^*-*1^, and commercial index (CI).

**Figure S4.**
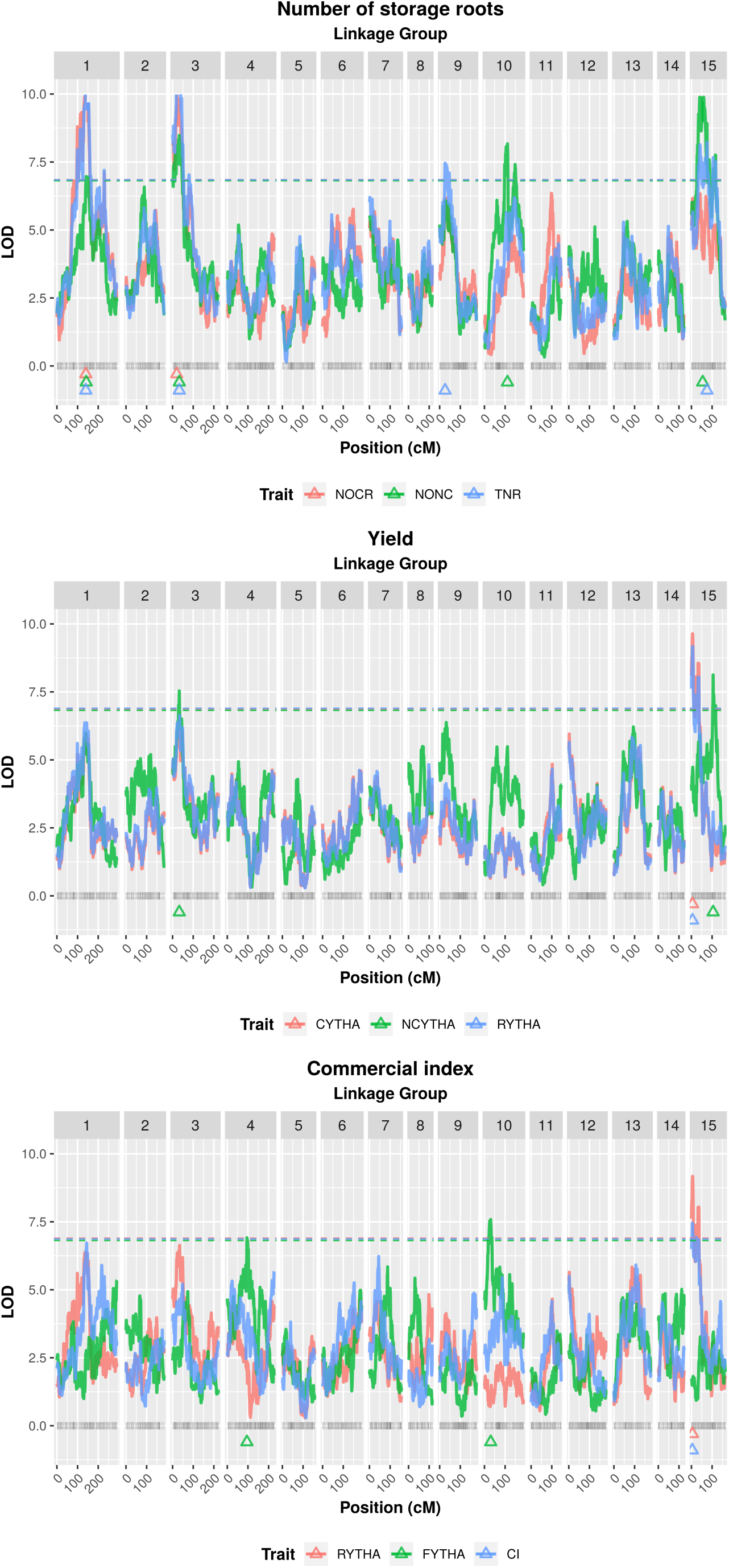
Logarithm of the odds (LOD score) profiles from fixed-effect interval mapping (FEIM) of eight yield-related traits from ‘Beauregard’ × ‘Tanzania’ (BT) full-sib family. Triangles represent the QTL peaks. Trait abbreviations: number of commercial (NOCR), noncommercial (NONC) and total (TNR) roots per plant, commercial (CYTHA), noncommercial (NCYTHA) and total (RYTHA) root yield in t · ha^*-*1^, foliage yield (FYTHA) in t · ha^*-*1^, and commercial index (CI). Dashed horizontal lines represent the permutation-based genome-wide significance LOD threshold of 0.20.

**Figure S5.**
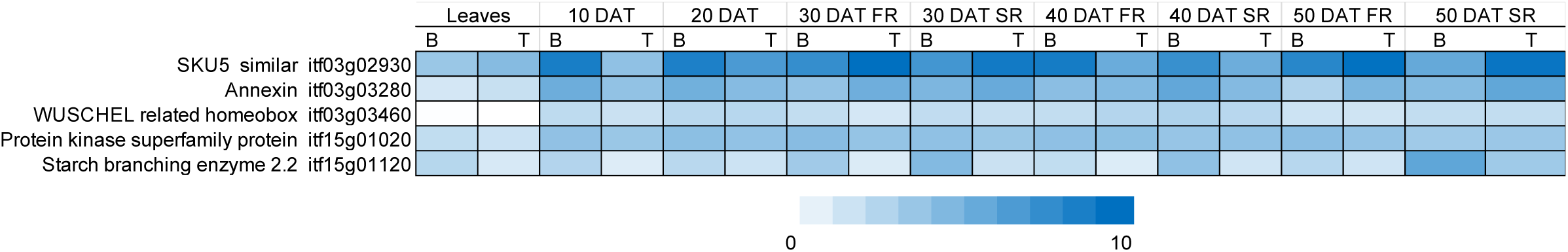
Heatmap with expression abundances in fragments per kilobase exon model per million mapped reads (FPKM, log_2_- transformed) for five genes in leaves and roots of ‘Beauregard’ (B) and ‘Tanzania’ (T). DAT: days after transplanting. SR: storage roots. FR: fibrous roots.

## Supplemental Tables

**Table S1.** Detection power (in percentage) and absolute difference between simulated and mapped QTL peak position (on average, in centiMorgans) from 1,000 simulated quantitative traits with three QTLs with different heritabilities 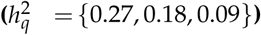. Fixed-effect interval mapping (FEIM) and random-effect multiple interval mapping (REMIM) were carried out under different genome-wide significance LOD and pointwise backward *p*-value thresholds, respectively, in ‘Beauregard’ × ‘Tanzania’ (BT) full-sib population

| $h_q^2$ | FEIM | | | REMIM | | |
| --- | --- | --- | --- | --- | --- | --- |
|  | Genome-wide significance | Power (%) | Difference (cM)<sup>a</sup> | Pointwise significance | Power (%) | Difference (cM)<sup>a</sup> |
| 0.27 | 0.20 | 89.40 | 2.41 (3.31) | $10^{-2}$ | 92.60 | 2.75 (3.37) |
| 0.18 |  | 73.00 | 3.16 (3.67) |  | 84.90 | 3.51 (3.93) |
| 0.09 |  | 34.50 | 4.06 (4.19) |  | 62.50 | 4.12 (4.13) |
| 0.27 | 0.15 | 88.80 | 2.40 (3.31) | $10^{-3}$ | 92.00 | 2.65 (3.36) |
| 0.18 |  | 71.60 | 3.16 (3.67) |  | 81.80 | 3.29 (3.81) |
| 0.09 |  | 32.30 | 3.99 (4.21) |  | 51.70 | 3.77 (3.97) |
| 0.27 | 0.10 | 88.00 | 2.40 (3.32) | $10^{-4}$ | 89.10 | 2.48 (3.24) |
| 0.18 |  | 70.00 | 3.16 (3.69) |  | 73.40 | 3.14 (3.73) |
| 0.09 |  | 29.90 | 3.85 (4.08) |  | 39.30 | 3.67 (3.89) |
| 0.27 | 0.05 | 86.60 | 2.38 (3.33) | $10^{-5}$ | 84.40 | 2.38 (3.17) |
| 0.18 |  | 66.80 | 3.13 (3.68) |  | 65.60 | 2.89 (3.59) |
| 0.09 |  | 27.20 | 3.82 (4.07) |  | 26.70 | 3.43 (3.86) |

<sup>a</sup> Standard errors of means are between parentheses.

**Table S2.** False discovery rate (FDR, in percentage) and proportion of matched QTL (coverage, in percentage) relative to support intervals calculated for each *d* = {1.0, 1.5, 2.0} from 1,000 simulated quantitative traits. Fixed-effect interval mapping (FEIM) and random-effect multiple interval mapping (REMIM) was carried out under different genome-wide significance LOD and pointwise backward *p*-value thresholds, respectively, in ‘Beauregard’ *×* ‘Tanzania’ (BT) full-sib population

| $d$ | FEIM | | | REMIM | | |
| --- | --- | --- | --- | --- | --- | --- |
|  | Genome-wide significance | FDR (%) | Coverage (%) | Pointwise significance | FDR (%) | Coverage (%) |
| 1.0 | 0.20 | 27.49 | 85.98 | $10^{-2}$ | 48.95 | 90.21 |
| 1.5 |  | 20.53 | 94.51 |  | 46.03 | 95.67 |
| 2.0 |  | 17.53 | 98.32 |  | 45.25 | 97.08 |
| 1.0 | 0.15 | 25.98 | 85.99 | $10^{-3}$ | 26.23 | 91.26 |
| 1.5 |  | 18.75 | 94.50 |  | 22.77 | 95.79 |
| 2.0 |  | 15.66 | 98.34 |  | 21.71 | 97.16 |
| 1.0 | 0.10 | 24.01 | 86.00 | $10^{-4}$ | 16.73 | 91.82 |
| 1.5 |  | 16.61 | 94.52 |  | 12.96 | 95.94 |
| 2.0 |  | 13.50 | 98.30 |  | 11.92 | 97.13 |
| 1.0 | 0.05 | 21.71 | 86.08 | $10^{-5}$ | 12.90 | 92.36 |
| 1.5 |  | 14.25 | 94.49 |  | 9.56 | 95.87 |
| 2.0 |  | 10.97 | 98.27 |  | 8.62 | 97.00 |

**Table S3.** Summary of fixed-effect interval mapping (FEIM) for eight yield-related traits in ‘Beauregard’ × ‘Tanzania’ (BT) full-sib family. Linkage group (LG), map position (in cM) and its *∼*95% support interval (SI), likelihood-ratio test (LRT), its corresponding logarithm of the odds (LOD) and the adjusted *R*^2^ (in percentage) are shown for each mapped QTL using permutation-based genome-wide significance LOD threshold of 0.20.

| Trait<sup>a</sup> | QTL | LG | Position (SI) | LRT | LOD | $R^2_{\text{adj}}$ (%) |
| --- | --- | --- | --- | --- | --- | --- |
| NOCR | 1 | 1 | 140.03 (113.05-154.02) | 47.71 | 10.36 | 11.83 |
|  | 2 | 3 | 20.18 (16.60-39.52) | 47.41 | 10.3 | 11.74 |
| NONC | 1 | 1 | 142.10 (127.44-162.44) | 32.08 | 6.97 | 7.08 |
|  | 2 | 3 | 32.56 (4.68-50.15) | 39.02 | 8.47 | 9.22 |
|  | 3 | 10 | 111.40 (97.02-154.13) | 37.61 | 8.17 | 8.79 |
|  | 4 | 15 | 54.49 (34.37-75.28) | 47.48 | 10.31 | 11.76 |
| TNR | 1 | 1 | 140.03 (131.50-155.34) | 49.28 | 10.7 | 12.29 |
|  | 2 | 3 | 32.56 (18.43-44.03) | 50.48 | 10.96 | 12.64 |
|  | 3 | 9 | 26.14 (15.11-54.08) | 34.35 | 7.46 | 7.78 |
|  | 4 | 15 | 75.28 (33.13-118.00) | 37.88 | 8.23 | 8.87 |
| CYTHA | 1 | 15 | 5.27 (1.07-36.02) | 44.4 | 9.64 | 10.84 |
| NCYTHA | 1 | 3 | 32.56 (24.47-44.03) | 34.71 | 7.54 | 7.89 |
|  | 2 | 15 | 105.02 (103.05-117.01) | 37.43 | 8.13 | 8.73 |
| RYTHA | 1 | 15 | 5.27 (1.07-36.02) | 42.23 | 9.17 | 10.19 |
| FYTHA | 1 | 4 | 95.01 (68.09-112.02) | 31.84 | 6.91 | 7.00 |
|  | 2 | 10 | 30.31 (16.12-37.32) | 34.93 | 7.58 | 7.96 |
| CI | 1 | 15 | 4.19 (0.00-37.25) | 34.31 | 7.45 | 7.77 |

<sup>a</sup> Trait abbreviations: number of commercial (NOCR), noncommercial (NONC) and total (TNR) roots per plant, commercial (CYTHA), noncommercial (NCYTHA) and total (RYTHA) root yield in $\text{t} \cdot \text{ha}^{-1}$ , foliage yield (FYTHA) in $\text{t} \cdot \text{ha}^{-1}$ , and commercial index (CI).

**Table S4.** Allele additive effects from QTL mapped for eight yield-related traits in ‘Beauregard’ × ‘Tanzania’ (BT) full-sib family using random-effect multiple interval mapping (REMIM). ‘Beauregard’ ({ *a*, …, *f* } ) and ‘Tanzania’ ( {*g*, …, *l*} ) alleles represent the parental contribution to the trait mean.

| Trait<sup>a</sup> | QTL | ‘Beauregard’ |  |  |  |  |  | ‘Tanzania’ |  |  |  |  |  |
| --- | --- | --- | --- | --- | --- | --- | --- | --- | --- | --- | --- | --- | --- |
|  |  | *a* | *b* | *c* | *d* | *e* | *f* | *g* | *h* | *i* | *j* | *k* | *l* |
| NOCR | 1 | -0.0767 | 0.0998 | -0.0025 | 0.0248 | -0.0221 | -0.0233 | -0.0168 | 0.0216 | 0.0305 | 0.0523 | -0.1057 | 0.0181 |
|  | 2 | 0.0072 | 0.0782 | -0.0600 | 0.0351 | -0.0162 | -0.0444 | -0.0083 | 0.0966 | -0.0420 | 0.0219 | -0.0541 | -0.0140 |
|  | 3 | 0.0112 | -0.0360 | -0.1171 | 0.0791 | 0.0172 | 0.0456 | -0.1475 | 0.0045 | 0.0302 | 0.0685 | 0.0272 | 0.0171 |
|  | 4 | -0.0446 | -0.0138 | 0.0545 | -0.0276 | 0.0031 | 0.0285 | 0.0407 | -0.0534 | 0.0618 | -0.0010 | -0.0569 | 0.0088 |
|  | 5 | 0.1042 | -0.0060 | -0.0487 | -0.0179 | -0.0287 | -0.0029 | 0.0072 | 0.0802 | -0.0385 | -0.0659 | 0.0086 | 0.0083 |
|  | 6 | -0.0184 | -0.0463 | 0.0944 | 0.0299 | -0.0200 | -0.0396 | 0.0406 | -0.0627 | -0.0084 | -0.0149 | -0.0005 | 0.0459 |
|  | 7 | -0.0979 | 0.0436 | 0.0795 | -0.0238 | 0.0053 | -0.0066 | -0.0073 | -0.0213 | -0.0033 | -0.0188 | 0.0254 | 0.0254 |
| NONC | 1 | -0.0065 | 0.0898 | -0.0631 | -0.0086 | 0.0004 | -0.0120 | -0.0094 | 0.0480 | 0.0079 | -0.0105 | -0.0073 | -0.0286 |
|  | 2 | 0.0492 | -0.0402 | -0.0619 | 0.0578 | -0.0399 | 0.0350 | -0.0476 | -0.0012 | -0.0319 | 0.0295 | -0.0109 | 0.0622 |
|  | 3 | -0.0601 | 0.0219 | -0.0284 | 0.0097 | -0.0053 | 0.0623 | -0.0135 | -0.0232 | 0.0733 | 0.0096 | 0.0048 | -0.0509 |
|  | 4 | -0.0147 | -0.0056 | 0.0199 | -0.0666 | 0.0535 | 0.0134 | 0.0182 | 0.0024 | -0.0133 | -0.0574 | 0.0566 | -0.0066 |
|  | 5 | -0.0355 | -0.0245 | 0.1207 | -0.0647 | -0.0168 | 0.0207 | 0.0324 | -0.0387 | 0.0096 | 0.0079 | 0.0051 | -0.0163 |
|  | 6 | -0.0041 | -0.0330 | -0.0358 | -0.0239 | 0.0225 | 0.0743 | -0.0605 | 0.0054 | 0.0790 | -0.0100 | 0.0174 | -0.0314 |
|  | 7 | -0.0375 | -0.0169 | -0.0522 | -0.0189 | 0.0433 | 0.0823 | 0.0026 | 0.0304 | -0.0097 | 0.0073 | 0.0098 | -0.0404 |
|  | 8 | 0.0480 | 0.0233 | -0.0488 | -0.0325 | 0.0034 | 0.0065 | -0.0718 | 0.0583 | 0.0063 | -0.0527 | 0.0385 | 0.0214 |
|  | 9 | -0.0236 | 0.0210 | 0.0394 | -0.0462 | -0.0337 | 0.0431 | 0.0222 | 0.0696 | -0.0431 | -0.0368 | 0.0138 | -0.0257 |
| TNR | 1 | -0.1089 | 0.1489 | 0.0388 | -0.0386 | -0.0364 | -0.0038 | -0.0366 | 0.0279 | -0.0089 | 0.1466 | -0.1538 | 0.0248 |
|  | 2 | 0.0674 | -0.0521 | -0.2164 | 0.1942 | -0.0873 | 0.0942 | -0.2958 | 0.0116 | -0.0017 | 0.1910 | 0.0015 | 0.0935 |
|  | 3 | -0.0910 | 0.0071 | -0.0011 | 0.0510 | -0.0154 | 0.0495 | 0.0869 | -0.0924 | 0.1659 | 0.0022 | -0.0802 | -0.0824 |
|  | 4 | -0.0169 | 0.0362 | 0.0188 | -0.0017 | -0.0564 | 0.0201 | -0.0109 | -0.0672 | 0.0224 | 0.0496 | -0.0830 | 0.0893 |
|  | 5 | 0.0093 | -0.0954 | 0.0393 | -0.0886 | 0.0793 | 0.0561 | 0.0286 | -0.0325 | 0.0097 | -0.0735 | 0.0481 | 0.0197 |
|  | 6 | -0.0923 | -0.0610 | 0.1614 | -0.0188 | -0.0513 | 0.0620 | 0.0677 | -0.0757 | -0.0191 | -0.0217 | -0.0048 | 0.0536 |
|  | 7 | 0.0630 | -0.0419 | -0.0103 | -0.0742 | -0.0033 | 0.0667 | -0.0921 | 0.0702 | 0.0999 | -0.0205 | 0.0585 | -0.1160 |
|  | 8 | 0.0531 | 0.0498 | -0.0964 | -0.0560 | 0.0354 | 0.0141 | -0.1263 | 0.0350 | -0.0019 | -0.0542 | 0.0885 | 0.0590 |
|  | 9 | -0.0183 | 0.0886 | -0.0302 | -0.0284 | 0.0634 | -0.0751 | 0.0805 | -0.1203 | 0.0930 | -0.0222 | -0.0296 | -0.0014 |
|  | 10 | -0.0322 | 0.0995 | 0.0176 | -0.0951 | -0.0842 | 0.0944 | -0.0457 | 0.1152 | 0.0623 | -0.0294 | -0.0839 | -0.0186 |
| CYTHA | 1 | 0.3022 | 0.8168 | -0.6094 | -0.0872 | -0.1778 | -0.2447 | 0.0310 | 0.2950 | 1.0456 | -0.3993 | -0.4669 | -0.5054 |
| NCYTHA | 1 | -0.0293 | 0.0510 | -0.0056 | -0.0203 | -0.0169 | 0.0212 | 0.0075 | 0.0358 | 0.0014 | 0.0273 | -0.0984 | 0.0264 |
|  | 2 | -0.0021 | 0.0178 | 0.0192 | -0.0118 | -0.0464 | 0.0233 | -0.0081 | -0.0665 | -0.0043 | 0.0406 | -0.0114 | 0.0497 |
|  | 3 | -0.0346 | 0.0214 | 0.0028 | -0.0366 | 0.0557 | -0.0086 | 0.0044 | 0.0093 | 0.0535 | 0.0021 | -0.0511 | -0.0181 |
|  | 4 | -0.0092 | -0.0401 | -0.0260 | 0.0362 | -0.0271 | 0.0662 | -0.0336 | -0.0396 | 0.0529 | 0.0296 | 0.0200 | -0.0294 |
|  | 5 | 0.0000 | 0.0234 | -0.0298 | -0.0756 | 0.0141 | 0.0679 | -0.0543 | 0.0586 | 0.0442 | 0.0067 | -0.0033 | -0.0520 |
| RYTHA | 1 | -0.0115 | -0.2356 | -0.3605 | 0.1004 | 0.4067 | 0.1005 | 0.4142 | -0.2283 | -0.5186 | -0.3287 | 0.3150 | 0.3464 |
|  | 2 | 0.3820 | 0.0903 | -0.4735 | 0.1308 | 0.0187 | -0.1483 | 0.3776 | -0.1423 | -0.7335 | -0.2565 | -0.0959 | 0.8506 |
|  | 3 | 0.3013 | 0.8584 | -0.6686 | 0.0513 | -0.3013 | -0.2410 | 0.0457 | 0.4922 | 1.1574 | -0.5636 | -0.4004 | -0.7314 |
| FYTHA | 1 | 0.5933 | 0.2377 | 0.0278 | -0.0235 | -0.0610 | -0.7743 | 0.3693 | 0.0962 | -0.5706 | -0.0128 | 0.3894 | -0.2715 |
|  | 2 | 0.3460 | 0.2159 | 0.0281 | -0.6872 | -0.0797 | 0.1769 | 0.6112 | -0.5162 | 0.0262 | 0.1795 | -0.3660 | 0.0653 |
|  | 3 | -0.2534 | -0.0520 | 0.0282 | 0.1661 | 0.2388 | -0.1277 | 0.5113 | -0.3626 | 0.0336 | 0.4025 | -0.0063 | -0.5785 |
|  | 4 | -0.2832 | -0.2848 | 0.2943 | 0.0672 | -0.1976 | 0.4041 | -0.2560 | -1.0853 | 0.6058 | 0.2293 | 0.4539 | 0.0524 |
|  | 5 | 0.1011 | 0.2950 | 0.0799 | 0.4343 | -0.7205 | -0.1898 | 0.0774 | -0.7765 | 0.3509 | 0.0161 | 0.0354 | 0.2968 |
| CI | 1 | 0.0012 | 0.0120 | -0.0060 | -0.0027 | 0.0004 | -0.0049 | 0.0012 | 0.0040 | 0.0104 | -0.0048 | -0.0056 | -0.0052 |

<sup>a</sup> Trait abbreviations: number of commercial (NOCR), noncommercial (NONC) and total (TNR) roots per plant, commercial (CYTHA), noncommercial (NCYTHA) and total (RYTHA) root yield in $\text{t} \cdot \text{ha}^{-1}$ , foliage yield (FYTHA) in $\text{t} \cdot \text{ha}^{-1}$ , and commercial index (CI).

**Table S5.** Map (in centiMorgans) and genome (in base pairs) positions and support intervals (inside the parentheses) for QTL mapped in ‘Beauregard’ × ‘Tanzania’ full-sib family using random-effect multiple interval mapping (REMIM) for eight yieldrelated traits.

| Trait<sup>a</sup> | QTL | LG | Map position (cM) | *I. trifida* genome (bp) | *I. triloba* genome (bp) |
| --- | --- | --- | --- | --- | --- |
| NOCR | 1 | 1 | 139.24 (99.06-152.48) | 22077567 (15917835-23281986) | 26692560 (10387193-28126436) |
|  | 2 | 2 | 139.30 (98.83-152.5) | 16126352 (11398424-3350738) | 11822379 (6161757-13945807) |
|  | 3 | 3 | 20.18 (0.00-37.44) | 1591872 (32030-3185578) | 1802096 (22131-3524655) |
|  | 4 | 4 | 76.48 (57.01-100.14) | 4780118 (3520761-7038218) | 5570824 (4107181-11896279) |
|  | 5 | 7 | 38.27 (2.18-99.12) | 2838315 (239482-15160871) | 3005905 (302031-21622505) |
|  | 6 | 9 | 31.15 (22.24-73.45) | 2036465 (1357520-3997473) | 2537129 (1804182-4799470) |
|  | 7 | 11 | 98.39 (83.07-123.22) | 7711459 (6002238-16415359) | 10309151 (8138260-18201378) |
| NONC | 1 | 2 | 88.62 (72.44-95.72) | 10660510 (9568479-11099215) | 5346836 (4151689-5850309) |
|  | 2 | 3 | 12.36 (0.00-21.01) | 790052 (32030-1645526) | 898334 (22131-1870158) |
|  | 3 | 4 | 65.09 (54.35-70.05) | 4039479 (3226706-4328193) | 4701624 (3769011-5022282) |
|  | 4 | 6 | 3.12 (0.00-25.13) | 10827111 (11507438-7099549) | 8590172 (9365714-4034190) |
|  | 5 | 9 | 15.11 (13.48-51.17) | 963597 (777490-2666908) | 1377719 (1161653-3269774) |
|  | 6 | 10 | 145.40 (137.16-164.81) | 22087839 (21754871-23272183) | 26130519 (25776790-27501919) |
|  | 7 | 12 | 137.16 (125.03-147.31) | 21521557 (20738759-22197168) | 25320356 (24372733-26179595) |
|  | 8 | 13 | 65.36 (50.68-98.41) | 12302173 (4360140-17295029) | 19828339 (8041557-25122550) |
|  | 9 | 15 | 46.19 (32.15-119.04) | 2779657 (1939509-18430098) | 3166284 (2238681-22171700) |
| TNR | 1 | 1 | 140.03 (130.13-148.23) | 22212737 (20422516-23040691) | 26810094 (6478813-27858718) |
|  | 2 | 3 | 20.18 (18.43-25.28) | 1591872 (1428660-1971958) | 1802096 (1628157-2218567) |
|  | 3 | 4 | 62.03 (43.43-86.11) | 3832284 (2567373-5837890) | 4469117 (2988152-9198140) |
|  | 4 | 4 | 200.09 (164.01-205.15) | 30146741 (27537111-30647356) | 33756387 (30687469-34246129) |
|  | 5 | 6 | 3.12 (0.00-14.33) | 10827111 (11507438-9128510) | 8590172 (9365714-6585862) |
|  | 6 | 9 | 73.45 (13.48-82.10) | 3997473 (777490-4928686) | 4799470 (1161653-6032732) |
|  | 7 | 10 | 149.31 (144.46-185.31) | 22370642 (22001105-24392989) | 26474885 (26021589-28734008) |
|  | 8 | 13 | 71.22 (44.23-120.57) | 13078300 (3404827-18987490) | 20738566 (6812279-27040045) |
|  | 9 | 14 | 66.19 (59.27-79.27) | 13866336 (6439913-15641815) | 12047246 (13374139-19755290) |
|  | 10 | 15 | 109.10 (65.09-118) | 11529245 (4456148-18121488) | 13540937 (4998050-21802867) |
| CYTHA | 1 | 15 | 5.27 (0.00-32.15) | 477772 (21822-1939509) | 570552 (56000-2238681) |
| NCYTHA | 1 | 1 | 131.50 (113.05-152.48) | 20574531 (19617725-23281986) | 6295866 (8560921-28126436) |
|  | 2 | 4 | 180.36 (43.43-215.18) | 28612876 (2567373-31472749) | 32007820 (2988152-35147251) |
|  | 3 | 8 | 0.00 (0.00-76.43) | 19730 (19730-5572337) | 67559 (67559-6976839) |
|  | 4 | 10 | 60.24 (35.15-114.14) | 5641312 (2981461-19734636) | 7167078 (3609708-23944890) |
|  | 5 | 15 | 105.02 (75.28-125.3) | 10277885 (5578641-18573061) | 11851576 (6377917-22348321) |
| RYTHA | 1 | 8 | 115.54 (40.19-115.54) | 15710677 (2729207-15710677) | 17988854 (3362265-17988854) |
|  | 2 | 13 | 128.02 (90.20-145.54) | 19774175 (16115515-20717332) | 27997761 (24485922-29160603) |
|  | 3 | 15 | 4.19 (0.00-8.02) | 452966 (21822-656169) | 541410 (56000-772513) |
| FYTHA | 1 | 4 | 63.08 (0.00-112.02) | 3949338 (42497-11940181) | 4583079 (27376-15353793) |
|  | 2 | 7 | 56.40 (34.12-109.48) | 4151772 (2632557-10523783) | 4391711 (2793744-15591910) |
|  | 3 | 8 | 43.00 (27.20-54.33) | 2990041 (2088002-3564360) | 3816702 (2602036-4509723) |
|  | 4 | 10 | 29.09 (18.40-37.32) | 2508929 (1725045-3121782) | 2930617 (1764312-3779221) |
|  | 5 | 13 | 42.60 (31.67-68.18) | 3275253 (2251209-12530096) | 6492727 (3912768-20079912) |
| CI | 1 | 15 | 5.27 (0.00-36.02) | 477772 (21822-2185604) | 570552 (56000-2490117) |

<sup>a</sup> Trait abbreviations: number of commercial (NOCR), noncommercial (NONC) and total (TNR) roots per plant, commercial (CYTHA), noncommercial (NCYTHA) and total (RYTHA) root yield in t · ha<sup>-1</sup>, foliage yield (FYTHA) in t · ha<sup>-1</sup>, and commercial index (CI).

